# The cellular and behavioral blueprints of chordate rheotaxis

**DOI:** 10.1101/2025.03.22.644710

**Authors:** Oleg Tolstenkov, Ayumi Ozawa, Célestine Allombert-Blaise, Jørgen Høyer, Tetsuya Hiraiwa, Marios Chatzigeorgiou

## Abstract

The oceans are filled with life-forms exhibiting complex adaptations to help navigate fluid environments and take advantage of fluid motion to locomote and disperse. However, the neuronal and behavioral underpinnings of navigating in fluid environments outside vertebrates are poorly understood.

We present behavioral and computational modelling-based evidence that the pre-vertebrate chordate *Ciona intestinalis* actively modulates its heading and angular velocities to counter oncoming flows and perform positive rheotaxis.

We demonstrate that a distributed network of ciliated peripheral sensory neurons is responsible for sensing hydrodynamic information such as flow velocity and direction. Pharmacological removal of sensory cilia impedes rheotactic behavior and reduces stimulus evoked neuronal activity.

Whole-brain calcium imaging experiments reveal that the central nervous system of *Ciona* can encode the onset and offset of flow, flow direction, and flow velocity, suggesting that miniature chordate brain is capable of surprisingly sophisticated computations.

## INTRODUCTION

The movement of organisms from one location to another has been a fundamental driving force of ecological and evolutionary processes since the beginning of life on Earth^1-3^. A broad range of species, which includes flying and swimming organisms, are almost continuously subjected to stronger or weaker flows of the air or water currents, respectively. Therefore, for animals that swim or fly, their movement is not only influenced by their locomotion but also by the motion of the fluid they are immersed in^4-6^. Consequently, animals that are involved in goal-oriented swimming or flying are faced with the requirement to utilize sensory and behavioral mechanisms that enable them to detect directional flows and correct their locomotion to profit from favorably directed flows or counter opposing flows^7^.

Paleontological records from the Ediacaran biota suggest that in aquatic environments, rheotaxis may have been one of the earliest mechanisms that evolved to exploit or counter turbulent flows^3,8-10^.

Rheotaxis is a form of oriented movement exhibited by an organism in response to a current of fluid^11-20^. Rheotactic behavior has been observed in organisms of very different dimensions, from millimeter long fish larvae^16,18^ all the way to meters long sharks^12^ including the enormous filter-feeding whale sharks (*Rhincodon typus*)^21^. The sensory basis of rheotaxis has been explored in fish using models like goldfish (*Carassius auratus)* and zebrafish (*Danio rerio*). Vision was regarded a key sense to detect whole-field motion relative to their frame of reference^11,22-25^ even though the idea of visual dominance has been challenged^26^. Indeed, aquatic organisms can perform rheotaxis in the absence of visual cues by leveraging mechanosensory information detected by the lateral line^13,16,18,27-30^. More recently, the idea of multisensory control of rheotaxis has been put forward^17,31^. Investigations of the circuitry architecture underlying flow sensing in fish have shown that lateral line ganglia respond to the direction of flow, while different brain regions encode flow onset, speed and accumulated displacement of flow^27,32^. Outside aquatic vertebrates rheotaxis remains an understudied behavior ^33-38^, while the evolution of the behavior and the underlying molecular and neuronal mechanisms remain to date unmapped.

To address these gaps in our knowledge we must turn our attention to marine invertebrate chordates that almost 500 million years ago, started to colonize the oceans, using their dispersing larvae which swam in the water column to reach new habitats. The ability to navigate in the water column and the capacity to hitchhike favorably directed flows or to counter opposing flows will likely have been critical to their ability to spread in the oceanic environment. Evidence of motility and associated behavior of extant invertebrate chordates are generally rare in the fossil record. Therefore, we must study extant invertebrate chordates like tunicates whose distribution in natural marine habitats has been attributed to water flow^39-41^.

However, the question of whether invertebrate chordates like the tunicates are capable of rheotaxis remains to date unanswered. More generally the behavioral algorithms that zooplanktonic organisms employ to perform rheotaxis and the underlying neuronal mechanisms are largely unknown. Importantly it is unclear how zooplanktonic larvae encode sensory cues (e.g. hydromechanical cues) in their brains to reach the adaptive decision to swim with or against the direction of the water flow. Here, we leverage the zooplanktonic swimming larva of the tunicate *Ciona intestinalis*, with an extremely streamlined nervous system composed of only 231 cells ^42^, to investigate the behavioral and neuronal bases of rheotaxis.

Using quantitative behavior analysis and mathematical modeling we find that *Ciona* larvae exhibit rheotaxis, using a distinct behavioral algorithm, based on modulating their heading and angular velocities. Using Ca^2+^ imaging coupled to directional flow stimulation we demonstrate that polymodal sensory neurons which are distributed across the surface of the larval trunk show directional selectivity to oncoming current. By combining whole-brain Ca^2+^ imaging and microfluidics we determine that the larval midbrain is encoding flow onset and offset, while the hindbrain is processing flow velocity information. By investigating both behavioral and neuronal responses, we provide new insights into the mechanisms underlying flow sensing in marine invertebrates.

## RESULTS

### *Ciona* larvae exhibit positive rheotaxis by modulating heading and angular velocities

To investigate the ability of *Ciona intestinalis* larvae to perform rheotaxis we designed a rheotaxis setup capable of exposing multiple *Ciona* larvae to flowing streams whose velocity could be adjusted (Figure 1A-C). The larvae were imaged under near-infrared light to avoid visible light interference^43-46^. Using both trajectory density analysis and 2D markerless pose estimation ^47^ (Figure 1D and E), we were able to track and quantify the larvae’s motion. When exposed to a range of flow stimulus strengths (1-3mm/s), we observed that *Ciona* larvae were either swept from the flow or they exhibited positive rheotaxis (Figure 1F and 1G; Figure S1E; Video S1). Quantification of larval swimming revealed that the mean rheotaxis probability (probability of upstream orientation) was significantly higher when animals were challenged with a 2mm/s (two body lengths/s) flow stream compared to static (0 mm/s) water (Figure 1H; Figure S1A, S1C, and S1G; S1F, and S1I for other flow strengths). This positive rheotaxis response was characterized by a significant increase in average heading velocity (Figure S1B, S1D, S1L, S1N, and S1J; Figure S2A), while angular velocity, a metric of rotational velocity about the center of mass of the larva was reduced when swimming longer distances suggesting more direct swimming (Figure S1M and Figure S2A), while the averaged mean angular velocity did not change (Figure S1K, S1O), indicating the larvae’s ability to effectively compensate for displacements induced by the flow of the water. To further characterize the phenotype, we quantified the density of actively swimming *Ciona* larvae body vectors orientation. Under no flow conditions we observed a uniform density across all orientations (Figure 1I, blue density). When we introduced a flow stimulus, we saw a prominent peak at 90° (upstream) and a smaller peak at 270° (downstream) suggesting that the larvae aligned primarily against and secondarily with the direction of flow (Figure 1I, red density). Close inspection of swimming trajectories suggested that larvae in static sea water performed mostly localized circular swimming akin to what has been shown previously for spontaneously swimming larvae^48^, in contrast to larvae exposed to a 2mm/s flow which moved upstream following a helical swimming pattern (Figure 1J).

**Figure 1.**
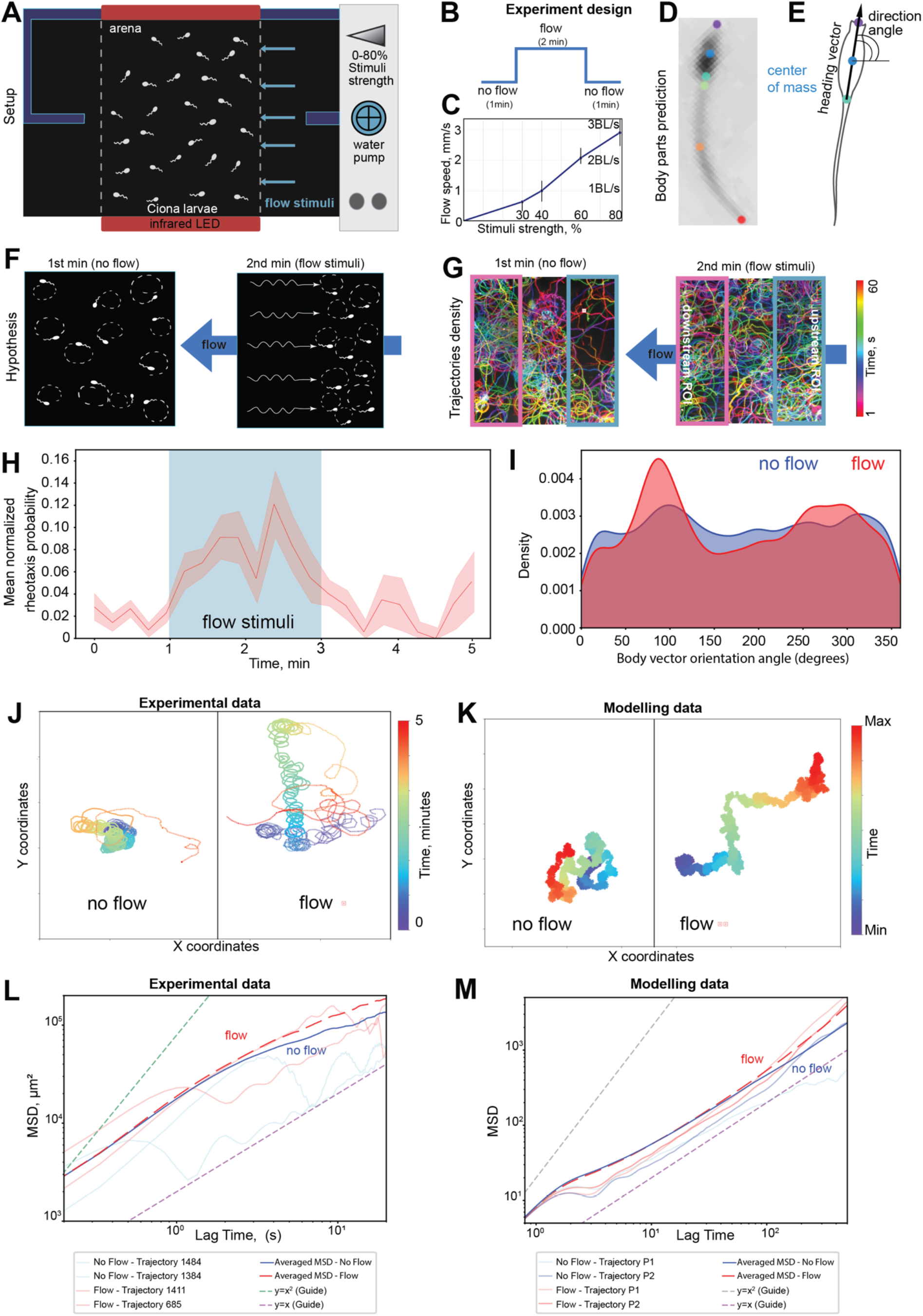
*Ciona* larvae exhibit positive rheotaxis by modulating heading and angular velocities. (**A**) Schematic of the rheotaxis setup. (**B**) Experimental protocol timeline: 1 min. of static water, followed by 2 min. of exposure to flow, followed by 2 min. of static flow. **(C)** A calibration plot showing the flow stimulus strength (% of maximum flow strength) as a function of water flow velocity in mm/s and *Ciona* body lengths (BL) per second. **(D)** A deep neural network (DNN) trained in DeepLabCut^47^ was used to extract the locations of *Ciona* body parts from videos. The image shows the most likely locations for 6 labeled body parts of a *Ciona* larva. **(E)** Schematic of the labelled body parts used for establishing the larva’s heading (body) vector and direction (orientation) angle between the heading vector and fixed coordinates of arena **(F)** Schematic illustrating the hypothesis of positive rheotaxis in swimming *Ciona* larvae. **(G)** The trajectory density of a representative experiment, alongside a schematic illustrating the measurement and comparison of mean grey values of animal trajectories in the upstream and downstream regions of interest (ROIs) under control (no flow) and flow stimulus conditions with 1 min duration for each condition. Trajectories are color-coded to represent different time points. **(H)** Mean rheotaxis probability (defined as the probability of upstream orientation) under no flow conditions and 60% strength, i.e. 2mm/s. Solid line shows the mean probability value, shade indicates SEM. Blue vertical bar defines the flow stimulus presentation for the flow experiments. **(I)** Body vector orientation distributions under no flow (blue) and 60% flow (red) conditions for wild-type larvae. Animal trajectories for H and I were filtered based on speed (>1.2 mm/s) to exclude inactive larvae (Supplementary Figure S1N). **(J)** Representative trajectories of swimming larvae exposed to no flow (static water) or flow at 2mm/s recorded in the rheotaxis setups (experimental data). **(K)** Representative trajectories which are the output of the mathematical model. For trajectories in J and K color-coding corresponds to temporal progression (blue to red). **(L and M)** Mean square displacement (MSD) based on experimental data (L) and computational simulation data (M) under no flow (solid blue curve) or flow stimulus (dotted red curve) conditions showing a shift (transition) of the movement pattern from a more diffusive regime (purple guideline) characterized by frequent reorientations to a more ballistic regime (green guideline) with more directed trajectories in the flow condition. The observed MSD decay at 10 s in both no-flow and flow conditions is likely due to the animal’s natural tendency to move in circular patterns.

To quantitatively investigate how a *Ciona* larva can achieve rheotaxis while executing a helical trajectory, we constructed a computational framework to simulate the dynamics of swimming *Cionas*. Here, we simply assume that, at the time *t*, a *Ciona* is located at the position (*x*(*t*), *y*(*t*)) with body vector orientation *θ*(*t*), and the equation of motion is given by

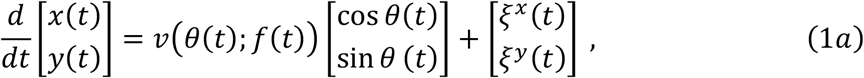

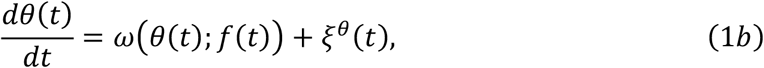

where *v*(*θ*; *f*) and *ω*(*θ*; *f*) represent the heading speed and angular velocity, which in general can depend on the body vector orientation *θ* and the flow condition *f*, the parameter indicating whether the flow stimulation is on or off; *f* = 1 (*f* = 0) in the presence (absence) of the flow. These functions can reflect the experimental observations. Note that, nicely, the mathematical analysis showed that *ω*(*θ*) should be proportional to 1/*h*(*θ*) (Supplementary Materials), which help us to set *ω*(*θ*) from the data. We assume that the system is exposed to noises, which are represented as *ξ*^*x*^(*t*), *ξ*^*y*^(*t*), and *ξ*^*θ*^(*t*) (see Supplementary Materials for more details).

Experimental results show that *v*(*θ*; *f*) and *ω*(*θ*; *f*) vary depending on the surrounding flow and the body vector orientation (Figure S4H). Specifically, the two peaks at approximately 90° and 270° in the body vector orientation distribution *h*(*θ*) under flow condition (Figure 1I) indicate that the larvae tend to change their orientation slowly when they direct these angles (Figure S2B). Furthermore, in the presence of flow stimulation, the heading speed of actively swimming larvae tends to be smaller when the body vector orientation *θ* is close to 90° and 270°. We incorporate in our model these responses to the flow by setting the functions *v*(*θ*; *f*) and *ω*(*θ*; *f*) to match to the experimental data (Figure S2B and S4H).

We confirmed that the computer simulations based on this model indeed yielded trajectories that resembled the experimental trajectories (Figure 1J and K, Figure S2C). Additionally, use of the mean square displacement (MSD) metric ^49^ suggested that experimental (Figure 1L) and model-derived data (Figure 1M) exhibit comparable movement dynamics in the following sense: (1) on a sufficiently short time scale, the slope of the MSD plotted on a log-log scale is approximately 1, indicating that the dynamics is diffusive, (2) on a larger timescale, the slope of the MSD under the flow condition is larger than 1. Thus, the MSD analysis supports the rheotaxis hypothesis, showing increased displacement and ballistic movement and indicating directed movement under flow conditions compared to more random, diffusive behavior in no-flow conditions (Figure 1L and 1M; Figure S1P).

These results indicate that the realization of the rheotaxis while drawing a helical trajectory can be explained by the modulation of heading speed and angular velocity depending on the relative angle of the body orientation to the flow direction and support the validity of our mathematical framework. To further resolve the contributions of the heading speed and angular velocity in achieving rheotaxis, we use the model simulated here as a control for further investigation under “one-sided” conditions. Namely, we compared the control with the following two cases: (1) the angular-velocity-only condition, where *ω*(*θ*; *f*) is the same as the control while *v*(*θ*; *f*) = *v*_0_, and (2) the heading-speed-only condition, where *v*(*θ*; *f*) is the same as the control while *ω*(*θ*; *f*) = *ω*_0_. The resultant MSD behaves similarly to the control when only the *ω*(*θ*; *f*) follows the case with flow, whereas it does not when only the heading speed follows the case with flow (Supplemental Material; Figure S2D-S2F). We thus conclude that, while both heading speed and angular velocity are modulated by flow, *Ciona* rheotaxis is caused specifically by the modulation of the angular velocity. While the principal mechanism of the rheotaxis is the modulation of the angular velocity, further numerical investigation by partly modifying *v*(*θ*; *f*) indicated that the modulation of the heading speed around *θ* = 270° enhances the rheotaxis. See Supplemental Materials for more details.

### A distributed network of polymodal sensory neurons detects changes in water flow

Despite the prevalence of rheotactic behavior in nature, little is known about the underlying neural mechanism outside a handful of model organisms^16,27,32^. We hypothesized that *Ciona* employs its miniaturized nervous system to perceive and encode changes in flow which result in rheotactic behavior. *Ciona* is equipped with 54 peripheral sensory neurons^50^, distributed across the trunk and tail of the larvae. These neurons are equipped with cilia or ciliary neurites that protrude into the tunic^51,52^. They function either as primary mechanosensory neurons^53^ or as polymodal sensory neurons capable of responding to mechanical and chemical cues important for larval settlement and metamorphosis^54-57^.

To test the ability of the sensory cells to detect water flow, we conducted experiments using a perfusion pencil-based flow system (Figure 2A) combined with calcium imaging in larvae expressing the calcium indicator GCaMP6s^58^ using the *Cii*.*pc2* promoter which labels robustly peptidergic neurons, which includes the majority of the peripheral nervous system and a subset of CNS neurons^54,59^ (Figure 2B and C and Figure S2).

**Figure 2.**
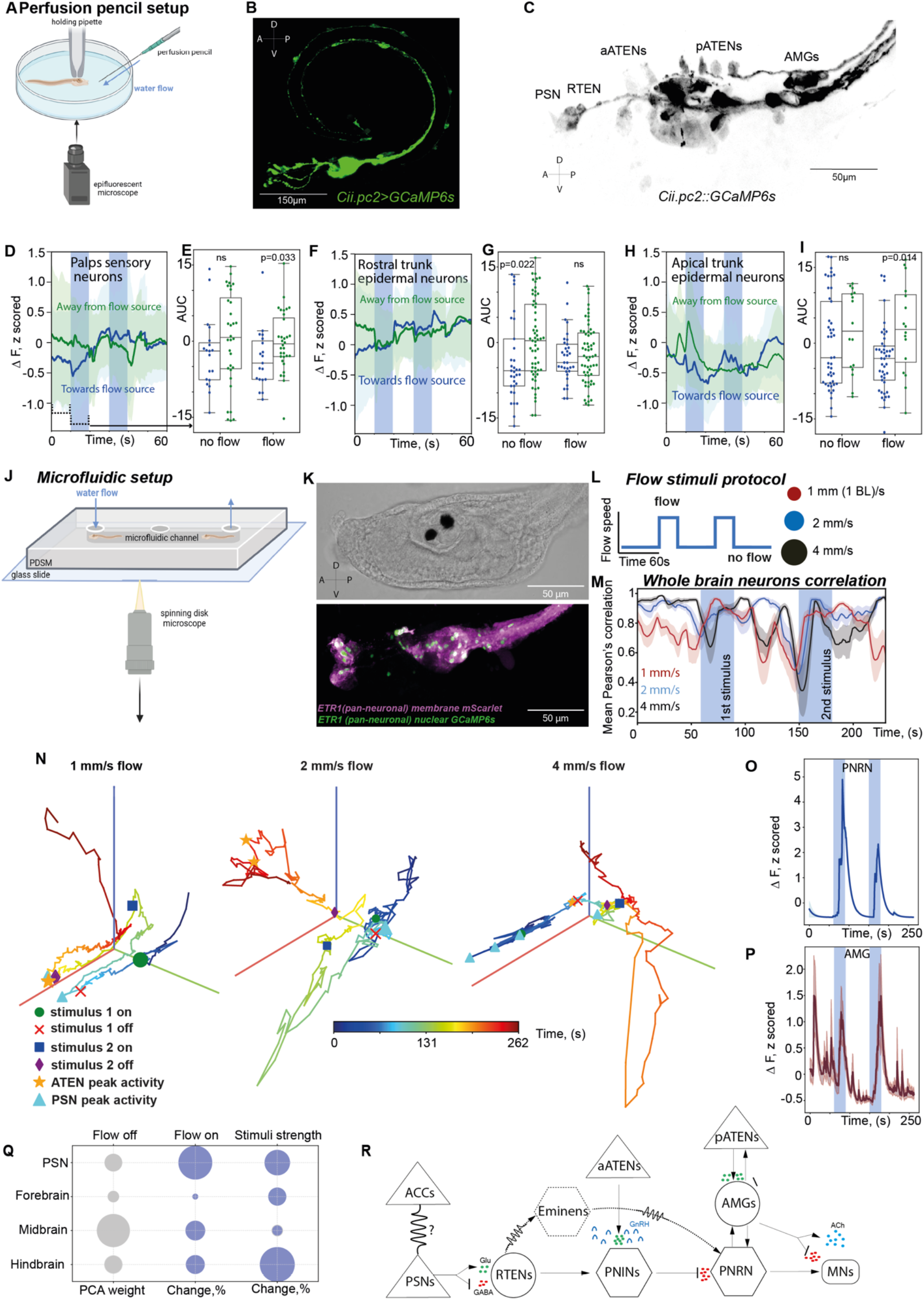
A distributed network of polymodal sensory neurons detects changes in water flow and engages midbrain and hindbrain neurons to compute flow features. (A) Schematic of the perfusion pencil setup used to deliver flow stimuli while performing Ca^2+^ imaging. (B) Representative maximal projection of a *Ciona* larva expressing *Cii*.*pc2::GCaMP6s* in peptidergic neurons of the CNS and Peripheral Nervous System (PNS) (C) Max projection of trunk neurons labelled by *Cii*.*pc2::GCaMP6s* in *Ciona* larvae, the polymodal papillae sensory neurons (PSNs), the rostral trunk epidermal neurons (RTENs), the anterior/ posterior apical trunk epidermal neurons (aATENs & pATENs) and the ascending motor ganglion interneurons (AMGs). (D-I) Mean Ca^2+^ responses and quantification of area under the curve (AUC) to directional flow stimuli, showing directionally selective activation patterns of selected sensory cell types when they are positioned towards or away from the flow source. The solid line indicates mean response; shaded areas show standart deviation SEM. Blue intervals indicate flow stimulation. Data points in (E,G,I) represent individual neurons. Dotted lines in D show the time intervals for mean quantative comparison. Statistical analysis in (E, G and I) was performed using the Mann-Whitney U test. (J) Schematic of the microfluidic setup used for whole-brain Ca^2+^ imaging. (K) Brightfield and maximal confocal projection of the trunk region of a representative transgenic *Ciona* larva which was used for whole-brain imaging expressing *Cii*.*Etr-1*>*nls::GCaMP6s::nls* (green) and *Cii*.*Etr-1*>*Lck-mScarlet* (magenda). (L) Flow stimuli protocol that was applied to *Ciona* larvae in the microfluidic chip. Two flow stimuli were delivered per animal. (M) Mean Pearson’s correlation of whole-brain activity calculated for medians of all cells with the 30 frames rolling window showing that repeated stimulus presentation enhances brain synchronization, shaded areas show 95% confidence interval. (N) Temporal progression of whole-brain Ca^2+^ activity trajectories at different flow speeds (1, 2, 4 mm/s). (O) Average trace of multiple Ca^2+^ responses from a class of premotor interneuron called peripheral relay neuron (PNRN). (blue bars indicate stimulus delivery interval). (P) Average Ca^2+^ responses from a class of peripheral motor neurons (AMGs). (Q) Quantitative summary of the impact of different brain regions on the processing of different parameters of flow stimuli. *Ciona* neurons were classified into brain regions as follows: PNS (PSNs, RTENs, ATENs, and DCENs), forebrain (PRs and coronet cells), midbrain (PNSRN, PRRN, ANTRN, and ANT), and hindbrain (AMG, PMG, and MN). PCA absolute loading weights were summed for each neuron type and aggregated by the brain region and time period. Changes in PCA weights from the no flow to flow condition and across different flow speeds are represented in the bubble plot. Grey bubbles indicate baseline activity; blue bubbles represent normalized changes relative to this baseline. (R) Putative neural circuitry transmitting flow cues from the periphery to the central nervous system and motor neurons, integrating sensory cell inputs and whole-brain imaging results. The signal propagates through the nervous system in accordance with predictions of the connectome. Glutamatergic/Gabaergic sensory cells (triangular cells) excite midbrain and hindbrain interneurons and relay neurons (polygons: Eminens, PNINs, PNRNs, circle: AMGs), which in turn balance excitatory and inhibitory signaling via acetylcholine (ACh) and GABA release to regulate the motorneurons (MNs, rectangle shape) of *Ciona*. Neurotransmitters and neuromodulators are assigned based on previous studies^74-80^.

Among the various types of sensory cells, the papillae sensory neurons (PSNs), the rostral trunk epidermal neurons (RTENs) and apical trunk epidermal neurons (ATENs) responded to flow (Figure 2D-I and Figure S3A to C). A feature of these Ca^2+^ responses is that their strength, as quantified by the area under the curve, showed a dependency on the flow direction (Figure 2D-I), suggesting that *Ciona* epidermal sensory neurons can encode the directionality of flow stimuli.

To investigate the downstream processing of flow stimuli, we performed whole-brain imaging of *Ciona* larvae using a microfluidics device (Figure 2J), with a nuclear localized GCaMP6s expressed pan-neuronally^54^ (Figure 2K). Interestingly, exposing larvae to two-flow stimuli protocol (Figure 2L) we found that all stimulus strengths trigger transient desynchronization, with the highest strength producing the most profound and most prolonged decrease in correlation. Recovery after the first stimulus is rapid in comparison to the second stimulus which causes a stronger disruption, suggesting cumulative effects or stimulus-induced adaptation. (Figure 2M). The speed of the flow cue with which the larvae were stimulated (we tested 1mm/s, 2mm/s and 4mm/s), was also encoded at a brain-wide level as it can be inferred by the qualitatively distinct brain activity trajectories elicited by the different flow speeds (Figure 2N). Downstream integrating premotor interneurons located in the midbrain (PNRN) and hindbrain (AMGs), played a major role in this processing (Figure 2O, 2P and Figure S3D). These neurons are highly relevant for integrating rheotactic stimuli information since the PNRN receives indirect inhibitory input from the RTENs and the AMGs receive direct input from the pATENs^42,50,60^.

To estimate the contribution of *Ciona*’s different brain regions in the stimuli information processing we quantified and grouped absolute PCA loading for all individual cells according to the anatomical region they were in (Figure 2Q). We determined that the midbrain was encoding flow onset and offset, while the hindbrain was processing flow speed information transmitted by sensory neurons to the CNS (Figure 2Q and R). Taken together our data suggest that *Ciona* senses flow using a distributed network of sensory neurons on the larval trunk and uses its midbrain and hindbrain circuitry to compute flow features like stimulus onset/offset and strength.

### Ciliated sensory neurons in the trunk and tail of the larvae are required for rheotaxis

In zebrafish, ablation of cilia on superficial neurons in the neuromast ridges significantly reduces rheotaxis ability ^16^. To test the role of ciliated sensory neurons in *Ciona* larvae, we chemically ablated the cilia using chloral hydrate ^61^ (Figure 3A-C). Loss of sensory cilia which protrude into the tunic resulted in a strong rheotaxis defect (Figure 3D, Video S2) including the reverse effect of the flow on heading and angular velocity compare to the wildtype (fig S4 A, B and D, E) and on the motion (fig S4 C). Upon the introduction of a flow stimulus control larvae performed a significantly higher number of upstream swims compared to downstream swims, thus exhibiting positive rheotaxis (Figure 3E and F). In contrast chloral hydrate treated animals did not significantly switch from downstream to upstream swimming upon flow introduction (Figure 3E and F). In addition, we quantified the density of *Ciona* larvae body vectors in chloral hydrate treated animals (Figure 3G). Chloral hydrate treated animals similarly to controls showed a uniform distribution of body vectors in static water (no flow) (Figure 1I; Figure 3G). However, in contrast to controls in the presence of flow we observed only the 270° peak suggesting that the larvae were unable to orient themselves against the flow (Figure 1I; Figure 3G and fig S1H, I, J, K). Finally, we observed a decrease in the probability to swim upstream upon introduction of flow (Figure 3H). We then decided to monitor Ca^2+^ responses to flow in the ciliated ATENs (located in the trunk of the animals) and the ciliated DCENs and VCENs located in tail. We found that chloride hydrate treatment eliminated flow induced Ca^2+^ responses in the ATENs but did not affect DCENs or VCENs responses (Figure 3I, 3J and Figure S4F and G). Taken together our results suggest that ciliated sensory neurons in the trunk of the animals are required for positive rheotaxis.

**Figure3.**
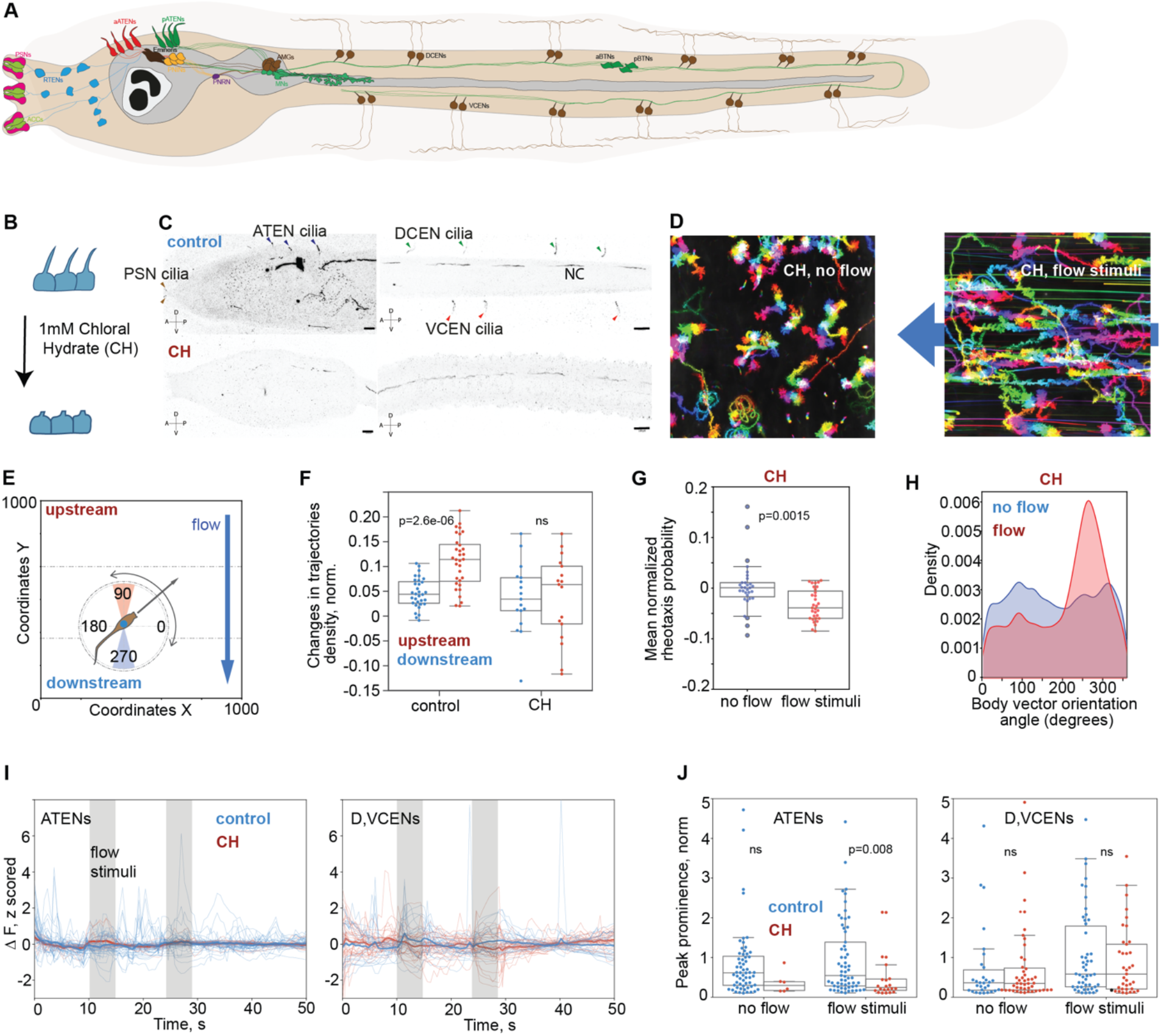
Trunk ciliated sensory neurons are required for positive rheotaxis. (A) Schematic of a Ciona larva showing key ciliated neurons of the PNS and some important CNS neurons. (B) Schematic of the chloral hydrate (CH) treatment used to ablate sensory cilia. (C) Maximal projections of control and chloral hydrate treated larvae where trunk and tail sensory neuron cilia, as well as ependymal cell cilia in the nerve cord are stained with anti-tubulin antibody. (D) Trajectories of chloral hydrate treated larvae under no flow and flow conditions for 1 min (color-code indicates temporal progression). (E) Schematic illustrating the range of angles considered as upstream swimming (red shade) and downstream swimming (blue shade). (F) Quantification of change in upstream and downstream trajectories density in wildtype and Chloral hydrate treated groups. Data points in F and H are representing the means of individual experiments. (G) Body vector orientation distributions under no flow (blue) and flow (red) conditions for chloral hydrate treated larvae showing the loss of the upstream angles peak in the treated animals. (H) Quantification of the probability to perform positive rheotaxis under no flow and flow conditions for chloral hydrate treated larvae. (I) Ca^2+^ traces showing reduced response in response to flow stimuli, in apical trunk epidermal neurons (ATENs) following cilia ablation relative to negative controls, while tail sensory neurons (DCENs, VCENs) remained unaffected. (J) Quantification of normalized peak prominence of Ca^2+^ transients elicited by flow stimuli in ATENs and DCENs/VCENs of control and chloral hydrate treated larvae. Data points in J are representing the individual neurons. For panels F, G, J we used a Mann-Whitney U test for statistical analysis.

## DISCUSSION

This study provides a systems level understanding of a miniature nervous system in a zooplanktonic invertebrate chordate larva and how it operates in marine habitats with dynamic fluid flows.

We establish that *Ciona* larvae can perform positive rheotaxis using helical swimming. Our experiments showed that a relatively complex behavior can be executed by a larva with such a small nervous system by using a behavioral algorithm whereby Ciona larvae can extract both directional information and speed information simultaneously. Zebrafish use a different but equally simple behavioral algorithm to perform rheotaxis using flow velocity gradients^18^. The rheotactic behavioral regime of *Ciona* differs also from that of other zooplanktonic organisms such as calanoid copepods (*Eurytenora affinis*), which use a mix of frequent jumps and turbulent transport^38^. The ability to detect and respond to different flow strengths may confer to *Ciona* the ability to detect and perform fast maneuvers to escape from predators as is the case for planktonic copepods^62^. Notably, even if most of the hatched larvae are swept from the water flow, the ability of a significant fraction to perform active rheotaxis may carry a strong fitness benefit for *Ciona* because it would enable these small zooplanktonic organisms to retain the advantages of self-locomotion under varying strength and direction flow conditions. In doing so it would give *Ciona* the capacity to move to a suitable habitat for settlement or to control their distribution despite strong hydrodynamics, while maintaining the diffusive properties of their motion. The result would be an ability to regulate the dispersal of the population akin to what has been documented for calanoid copepods^38^. This behavioral trait would provide a selective advantage even when the difference in diffusivity is small^63^.

This study has elucidated the sensory mechanisms and downstream processing circuit underlying flow sensation. We found that *Ciona* leverages a network of ciliated trunk epidermal sensory neurons which are distributed along the rostral/caudal and mediolateral axes to sense flow. We establish that these superficial sensory neurons detect flow over the surface of trunk and have a suitable spatial distribution to signal the direction and strength of flow, thus contributing to the detection of local differences in flow across different parts of the trunk. Epidermal neurons form a unique outer body dendritic network in the ascidian larval tunic ^52^. Interestingly they express differentiation gene batteries that are conserved with those of lateral line hair cells ^64^. In addition, an ancestral relationship to the chordate PNS has been postulated on the grounds that these cells deliver overlapping input to the CNS and that they are distributed in an antero-posterior direction^65^. We and others have recently shown that these epidermal sensory neurons detect both mechanical and chemical cues ^54,55^. While lateral line hair cells in vertebrates were thought to be exclusively mechanosensory, a recent study raises the possibility that they are polymodal, since they are also capable of sensing chemical cues^66^. Taken together, our findings suggest that the sensory cells employed by vertebrates to sense flow may have evolved earlier in the chordate lineage.

Through whole brain-calcium imaging we have identified at least two sensorimotor integration sites in the CNS, namely the PNRN neuron located in the midbrain and the AMGs located on the dorsal side of the hindbrain. Interestingly, the zebrafish hindbrain has been identified as the brain location where water-flow direction selective cell clusters are located^27,32^. In goldfish, midbrain and hindbrain clusters responded to various hydrodynamic stimuli^67,68^. We find that the peripheral nervous system and the hindbrain are two primary sites where flow strength is encoded and decoded in the *Ciona* larval nervous system. Early studies based on anatomical features, suggested that the peripheral DCENs and VCENs are initiating swimming or acting as proprioceptors during swimming^69^. While the DCENs may not contribute to sensing flow stimuli per se, they may provide tail postural information to their postsynaptic targets the AMG neurons, suggesting that the AMGs are key integrators of rheotactic information and self-movement.

Typically, animal brains including that of *Ciona* larvae transition between dynamic internal states in response to tasks such as foraging or chemosensation^54,70-73^. We show that *Ciona* has two distinct phases when engaging and disengaging in flow sensing, suggesting that state-encoding neural populations may be responsible for the initiation and termination of the rheotactic behavior. The emerging picture is that invertebrate chordates and teleost fish employ evolutionary conserved brain regions and neural mechanisms for sensorimotor transformation during rheotaxis.

## RESOURCE AVAILABILITY

### Lead contact

Further information and requests for resources and reagents should be directed to and will be fulfilled by the lead contact, Marios Chatzigeorgiou

### Materials availability

Plasmids generated in this study are available upon request from the lead contact.

### Data and code availability

- Raw calcium imaging data will be deposited in Zenodo and will publicly available as of the date of publication.
- All original code will be deposited at Zenodo and will be publicly available as of the date of publication. Code to analyze behavioral data, brain imaging data, 3D printing files are already available here: https://github.com/ChatzigeorgiouGroup/Rheotaxis_ciona_larvae
- Any additional information required to reanalyze the data reported in this paper will be made available from the lead contact upon request.

## ACKNOWLEDGMENTS

We would like to thank members of the Chatzigeorgiou lab for valuable feedback on the manuscript. We acknowledge funding from the Research Council of Norway: 339399 (M.C.) and 234817 to the Michael Sars Centre. JAMSTEC Young Research Fellow Research Grant (A.O.)

## AUTHOR CONTRIBUTIONS

Conceptualization: O.T., M.C.; Methodology: O.T., M.C., A.O., T.H.; Data curation: O.T., A.O., C.A.B., J.H.; Investigation: O.T., A.O., C.A.B., J.H.; Visualization: O.T., A.O., M.C., C.A.B., J.H.; Resources: O.T., M.C.; Funding acquisition: M.C., A.O.; Project administration: M.C., T.H.; Supervision: M.C.; Writing – original draft: O.T., M.C.; Writing – review and editing: O.T., A.O., M.C., C.A.B., J.H., T.H.

## SUPPLEMENTAL INFORMATION

**Document S1. Figures S1-S4**

**Movie S1. Rheotaxis response of wild type *Ciona intestinalis* larvae. Related to Figure 1**. Video of wild type *Ciona* larvae recorded in the rheotaxis behavioral setup, illuminated using infra-red light. The larvae respond to the 2 mm/s flow stimuli (from second to third minute of the video). Arrow shows direction of the flow.

**Movie S2. Loss of sensory cilia affects rheotaxis behavior of *Ciona intestinalis* larvae. Related to Figure 3**. Video of *Ciona* larvae treated with 5μM Chloral Hydrate (for 2 hours), which induces sensory cilia loss. The *Ciona* larvae do not show directional response to the 2 mm/s flow stimuli (from second to third minute of the video). Arrow shows direction of the flow.

## STAR★METHODS

### KEY RESOURCES TABLE

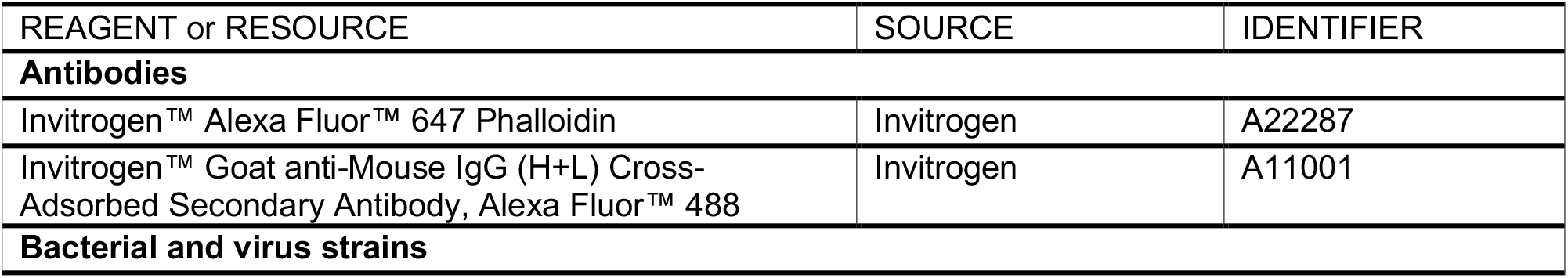
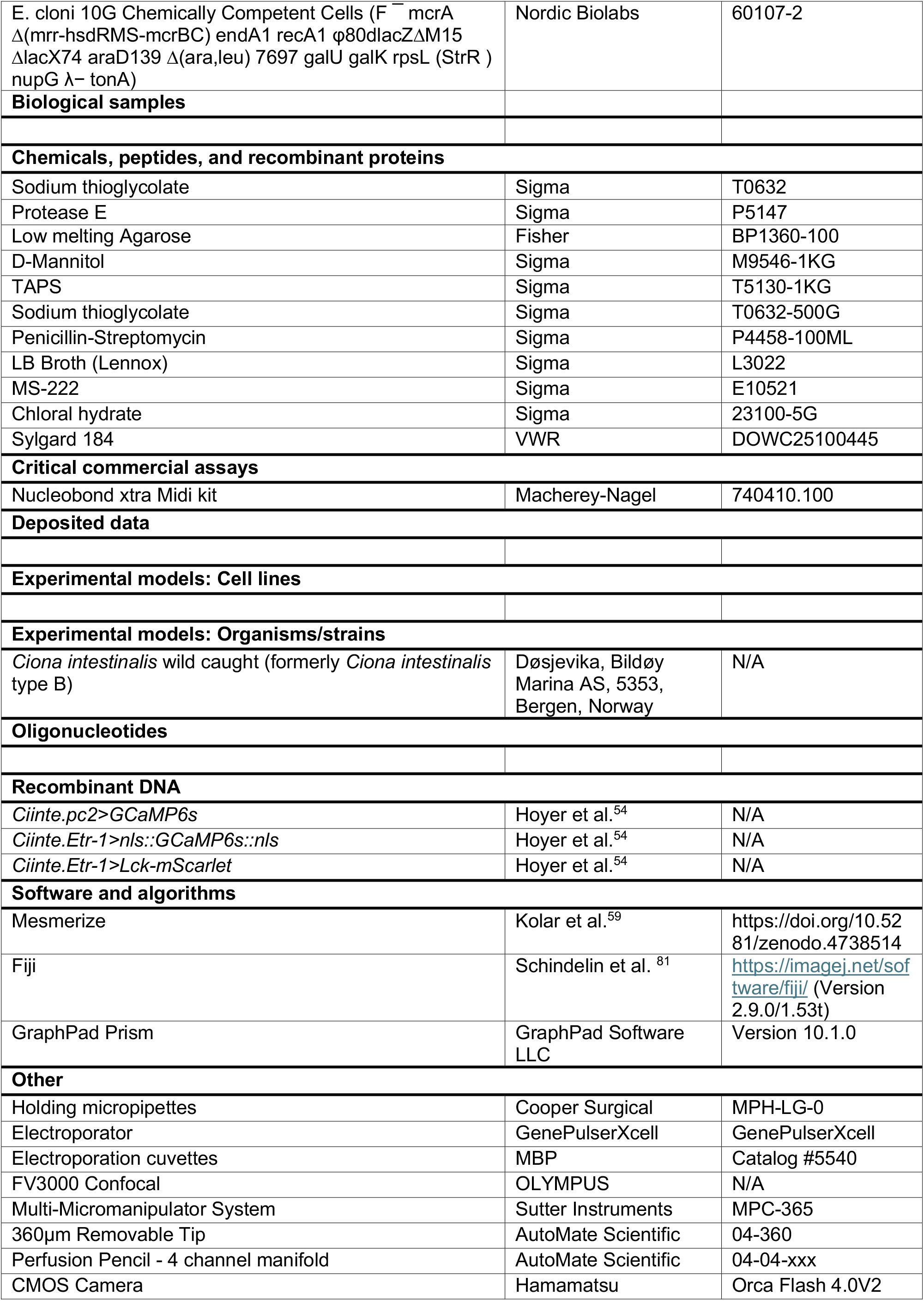
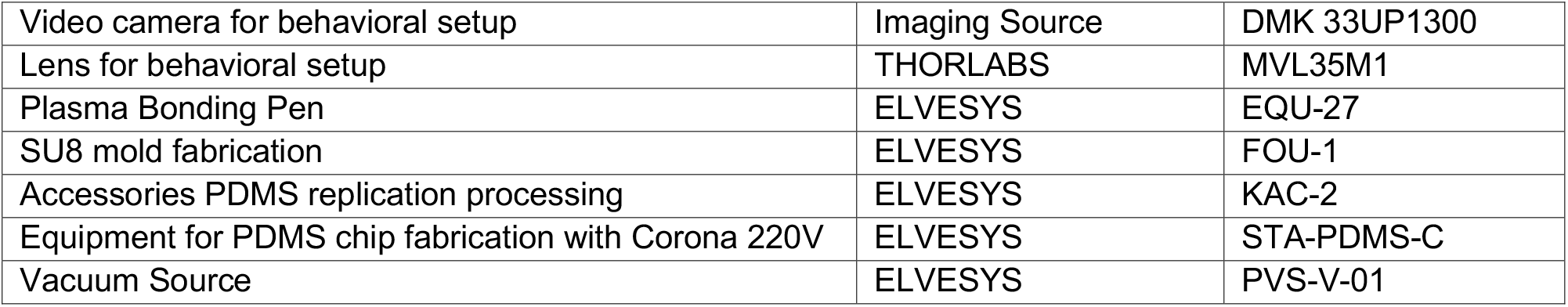

## EXPERIMENTAL MODEL AND STUDY PARTICIPANT DETAILS

We harvested adult *C. intestinalis* (i.e. *Ciona* intestinalis Type B) from: Døsjevika, Bildøy Marina AS, with postcode 5353, in Bergen, Norway. The collection site can be located using the GPS coordinates: 60.344330, 5.110812. We housed gravid adult *Cionas* in our purpose-built facility at the Michael Sars Centre. We kept approximately 100 adults in 50L tanks with constant running sea water at 10°C and provided constant light and food supply. The diet included several species of diatoms and brown algae to increase egg production and reduce spontaneous spawning^82^.

## METHOD DETAILS

### Electroporations of *Ciona intestinalis* zygotes

To obtain mature *C. intestinalis* eggs and sperm for *in vitro* fertilization we dissected adult *C. intestinalis*. To dechorionate *Ciona* eggs we carried out chemical dechorionation in a 1% sodium thioglycolate and 0.1% pronase mix dissolved in artificial filtered seawater (ASW). We provided mechanical dechorionation by rocking the eggs in pronase solution on a shaker for almost 10 minutes until we noticed that eggs were dechorionated. Using ASW we washed dechorionated eggs five times and then we fertilized with sperm diluted in ASW and Tris buffer for ∼10 minutes. After the successful completion of the fertilization we thoroughly washed the eggs from the residual sperm for 5 or 6 times with ASW. We carried out electroporations according to our previously published protocols with minor modifications^83^. The electroporation mixes were prepared by adding 400μl of 0.95M D-Mannitol and up to 100μl of DNA with 90 to 180 μg of DNA according to the expected levels of expression for each of the construct used. We electroporated the fertilized eggs using a 4 mm gap electroporation cuvette (MBP Catalog #5540) in which we transferred the electroporation mix and the fertilized eggs in 300μl ASW. To deliver the electrical pulses we used a BIORAD GenePulserXcell which had a CE-module. We used the following settings: Exponential Protocol: 50 V, Capacitance: between 1050 and 1300 μF, Resistance was set to: ∞. We obtained electroporation time constants between 10–25 milliseconds. We transferred electroporated zygotes in ASW which contained Pennicilin-Streptomycin antibiotic solution, at 14°C. Embryos were washed again the next day. We adjusted the pH of the ASW to 8.4 at 14°C. The salinity of the ASW was adjusted to 3.3–3.4%. The old ASW on the plates was replaced with fresh ASW and Pennicilin-Streptomycin antibiotics.

### Electroporated constructs and their cloning

For calcium imaging, three expression constructs were cloned as described in a recent publication^54^. We used *Ciinte*.*Etr-1*>*nls::GCaMP6s::nls* (150 μg of this construct to each electroporation mix) in combination with *Ciinte*.*Etr-1*>*Lck-mScarlet* (90 μg of this construct in each electroporation mix) for whole-brain imaging. For epifluorescence calcium imaging combined with stimulation with perfusion pencil, we used *Ciinte*.*pc2*>*GCaMP6s* construct of which we typically electroporated 150 to 180μg.

### Calcium imaging combined with perfusion pencil system

We imaged animals using a 40x water immersion objective mounted on an upright Axioskop A1 fitted with a Hamamatsu Orca Flash 4.0V2 CMOS camera. The usable field of view was 320μm in the x and y directions (2048×2048pixels). For illumination we used a mercury lamp and the following filter set: BP470/20, FT493, BP505-530. We acquired the data using the imaging software HCImageLive at 100Hz. The transient transgenic *Ciinte*.*pc2*>*GCaMP6s* larvae used for imaging were between 38 h.p.f. and 44 h.p.f. old (raised at 14°C). Larvae were transferred to the microscope and immobilized as described in Hoyer et al^54^. Larvae were oriented in the different positions with the holding pipette ^54^ relative to the perfusion pencil. We recorded an animal for 10 s before delivering 10 s of continuous flow of ASW.

### Whole brain imaging

For whole brain imaging we imaged using a 40x silicon oil objective, on an Olympus iXplore SpinSR10 spinning disk confocal equipped with two Hamamatsu Orca Flash 4.0V2 CMOS cameras. We cropped the FOV (1024×1024 with 2×2 pixel binning) to visualize only the trunk of the larva which was trapped in the microfluidic device. We illuminated the transgenic larvae using a 488nm laser. On average we acquired 356 volumetric cycles per movie. We adjusted the number of volumes we acquired to set the time of the experiment to 4 minutes. We used a step of 0.32μm on the z-axis within a range of 28.8μm. The data was acquired at approximately 1.5 brain volumes per second. The animals were firstly recorded without flow for 60 seconds before switching to a flow of 300, 150 and 75 mbar pressure for 30 seconds. We then repeated the 60 s no flow and 30 s flow cycle. To acquire the two-channel reference stacks we imaged both the nucleus and the plasma membrane (Lck-mScarlet marker), we illuminated the animal trunk using the 488nm and 561nm lasers with 0.5 μm step and 100 ms exposure time.

### Microfluidics chip construction and animal mounting

Electroporated embryos were raised in ASW at 14°C until hatching. The transient transgenic larvae used for imaging (between 38 h.p.f. and 44 h.p.f. old at 14°C) were sorted in 6-well plates that were coated with 1% agarose in ASW. The glass photomask was designed using KLayout software, following the design described by Guillaume Poncelet ^55^. The master mold in SU-8 was fabricated by ELVESYS, France for the channel height 42 and 35 µm. Microfluidic chips were produced using a prepolymer mixture of polydimethylsiloxane (PDMS, Sylgard 184 silicone elastomer kit, a complete kit of accessories for PDMS chip replication, KIT-APDMS). The mixture was prepared with a 1:10 ratio of curing agent to base polymer and cured at 65°C for 4 hours. The cured PDMS blocks were irreversibly bonded to 1.5H cover glasses using a Plasma Bonding Pen (EQU-27) with a 2-minute plasma treatment.

### Rheotaxis behavioural arena and experimental procedure

Chorionated eggs were fertilized with sperm for 10 minutes in the presence of 50μl

Tris buffer pH 9.5. Fertilized eggs in ASW were spread in multiple 9cm agarose coated dishes at 14°C until hatching (approximately 36-38h.p.f.). The swimming larvae were collected using a glass pipette under a binocular microscope and transferred to new 9cm agarose coated plates with the aim of concentrating and cleaning them from debris (e.g. unhatched eggs). 10 ml of larvae were transferred to the rheotaxis setup which was made from acrylic sheets and mesh that were glued together. The video recording was done with a DMK 33UP1300 (Imaging Source) coupled to a MVL35M1 (35mm EFL, F/1.4) lens. The rheotaxis chamber was made from acrylic plastic with the central chamber where we placed the animals having the following dimensions (width 25 mm, length 30 mm). Infrared illumination was provided by 8 IR (850nm wavelength) lights on either side of the arena. Flow was established using a DC12V pump with a maximum of 5000rpm and a maximum flow rate of 100 mL/min. The electronics of the setup were controlled via an Arduino and custom-made software written in Python.

The pre-stimulus period was 1 minute. The rheotactic stimulus was delivered for 2 minutes. The recording ended after 2 additional minutes. We calibrated the flow speed by measuring the mean of particles speed with the different pump flow rate.

### Chloral hydrate treatment and immunohistochemistry

Chorionated swimming larvae were incubated for 2 hours in 5 μM Chloral Hydrate (a 10 mM stock solution was diluted in ASW). After 2 hours the treated larvae were transferred to the arena to be assayed. A fraction of the treated larvae was fixed in MEM-FA buffer for 30 minutes at room temperature with nutation. To permeabilize the larvae, they were incubated in PBT buffer (0.15% Triton X-100 in PBS) for 5 minutes at room temperature. Subsequently, the larvae were incubated in PBTT-NH4Cl buffer, three times for 15 minutes at room temperature. The excess detergent was removed by washing the larvae three times for 10 minutes with TNT buffer. Then, the larvae were incubated in the blocking solution (TNBS) for 30 minutes at room temperature to block non-specific binding sites. Following this, the larvae were incubated with the primary antibody acetylated tubulin (Lys40) (Protein Tech, Cat. #66200-1-lg) diluted (1:500) overnight at 4°C. The excess primary antibody was removed by washing the larvae six times for 10 minutes with TNT buffer. To prepare the larvae for the secondary antibody, they were incubated in the blocking solution (TNBS) for 15 minutes at room temperature to block non-specific binding sites. Then, the larvae were incubated with the goat anti-mouse 488 fluorescently labelled secondary antibody (Invitrogen) diluted (1:500) in the blocking solution for 2 hours at room temperature in the dark. The excess secondary antibody was removed by washing the larvae six times for 10 minutes with TNT buffer. The stained larvae were mounted on slides and imaged using an OLYMPUS FV3000 confocal.

### Calcium imaging analysis

All video recordings were converted to TIFF stack time series and motion-corrected using the NoRMCorre algorithm ^84^. GCaMP6s intensity levels from imaged cells were extracted using the Mesmerize platform, employing CNMF-3D ^85,86^ for signal extraction in three-dimensional imaging. The extracted traces were z-scored, and principal component analysis (PCA) was performed for whole brain data to reduce dimensionality using the scikit-learn Python library ^87^. For visualization a three-component trajectory was obtained for each recorded sample. For quantitative comparison we used weights of five components. Cell identities were inferred by corroborating cell locations and neuronal morphologies—visualized using the *Ciinte*.*Etr-1*>*Lck-mScarlet* marker—with connectome data from Ciona intestinalis. This information was cross-referenced with the Cionectome-V2 dataset available on Morphonet (https://morphonet.org/dataset/2122)^88^.

### Theoretical models and numerical simulations

The computational model of a *Ciona* defined by Eq. (1a) and (1b) is based on the chiral active Brownian particle^89^, also known as the circle swimmer^90^. The chiral active Brownian particle is broadly used to model self-propelled objects that show circular trajectories and is given as below:

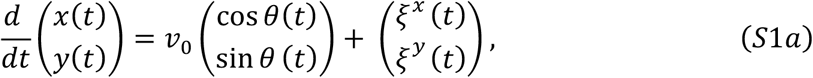

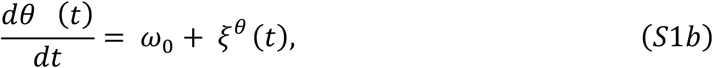

where *v*_0_ and *ω*_0_ are constants corresponding to the heading speed and angular velocity, respectively. We assume that the system is exposed to the white Gaussian noises, which are represented as *ξ*^x^(t), *ξ*^y^(t), and *ξ*^*θ*^(*t*). These random processes are assumed to be independent, i.e., ⟨*ξ*^*u*^(*t*)*ξ*^*v*^(*t*^′^)⟩ = *δ*_*u,v*_*δ*(*t* − *t*^′^)*D*_*u*_, where *u* and *v* are placeholders for *x, y*, and *θ, δ*_*u,v*_ is the Kronecker delta, *δ*(*t*) is the Dirac delta function, and *D*_*u*_ is the strength of the noise *ξ*^*u*^(*t*).

Equations (1a) and (1b) are obtained by replacing the constants *v*_0_ and *ω*_0_ in Eq. (S1a, S1b) with the functions *v*(*θ*; *f*) and *ω*(*θ*; *f*), respectively, where *f* is the parameter indicating whether flow stimulation is applied; *f* = 1 when the flow stimulation is on, and *f* = 0 otherwise. The functions *v*(*θ*; *f*) and *ω*(*θ*; *f*) were defined as follows based on the experimental data.

In the presence of the flow stimulation, the heading speed of *Ciona* tended to be smaller for *θ* close to 90° and 270°; see the red curve in Fig. S4(H). We incorporate this dependence by setting *v*(*θ*; 1) to be the mean of the instant heading speed over the animals with orientation angle *θ* under flow stimulation. More specifically, the range from 0 to 360°is divided into 72 bins with 5°intervals, and the mean heading speed 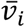 of the *i*th bin (*i* = 1,2, …, 72) is calculated by averaging the instant heading speed of animals whose orientation angle belongs to that bin. Thus, 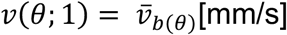 and *b*(*θ*) = ⌊72 *θ*/(2*π*)⌋, where ⌊ ⌋ denotes the floor function. In contrast, in the absence of flow stimulation, the heading speed did not show a clear dependence on the orientation angle (the blue curve in Fig. S4(H)). Hence, we simply set the heading speed to be constant for *f* = 0; *v*(*θ*; 0) = *v*_0_. The value of *v*_0_ is chosen to be the mean heading speed for 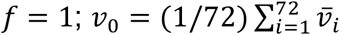.

The function *ω*(*θ*; *f*) was estimated from the distribution function of the angle *θ*. Note that, given the function *ω*(*θ*; *f*) > 0, the time evolution of the probability distribution function *g*(*θ, t*; *f*) of the angle *θ* is given by the following Fokker-Planck equation:

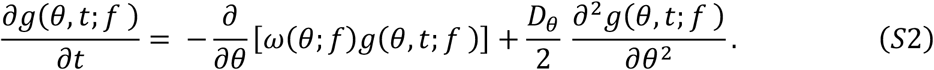

The steady solution ĥ(*θ*; *f*) of Eq. (S2) under the boundary condition ĥ(0; *f*) = ĥ(2*π*; *f*) and the normalization condition 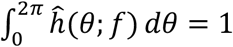 is given by^91^

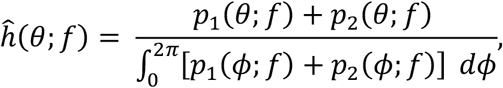

where

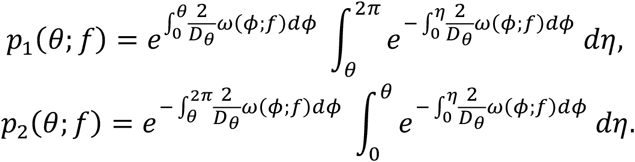

In the limit of *D*_*θ*_ → 0, ĥ(*θ*; *f*) approaches to *C*/*ω*(*θ*; *f*), where *C* is a constant that normalizes 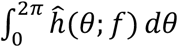 to be 1^91^. Hence, when the noise is assumed to be sufficiently small, *C*/ĥ(*θ*; *f*) gives the lowest-order approximation of *ω*(*θ*; *f*).

The experimental data indicated that the distribution of the body orientation angle has two troughs at approximately 90° and 270° under the flow condition, while the distribution in the absence of flow did not show clear dependence on the angle. Therefore, we set *ω*(*θ*; 0) = *ω*_0_ and *ω*(*θ*; 1) = *C*/*h*_*b*(*θ*)_, where *h*_*b*(*θ*)_ is the density of the *b*(*θ*)th bin under flow condition (Fig. S4(H)). The value of *ω*_0_ and *C* are chosen so that the mean 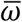 of *ω*(*θ*; *f*) with respect to *θ* s is close to the experimental value in the unit of [rad/s]. In Fig. 1K, 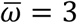 is used. To calculate the mean MSD in Fig. 1M, we used 100 particles whose 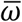 values were extracted from the uniform distribution in the interval from 0.6 to 5.4.

Note that *v*(*θ*; *f*) and *ω*(*θ*; *f*) can be multiplied respectively by arbitrary constants without loss of generality by rescaling space and time. From the mathematical point of view, this rescaling is equivalent to modifying the noise strengths *D*_*x*_, *D*_*y*_, and *D*_*θ*_. The values of *v*_0_ and *ω*i are chosen as above only for easy comparison with experimental data. The strengths of the noises are chosen to be sufficiently small so that the trajectory of the particle looks like those of *Ciona*, but sufficiently large so that a diffusive regime is observed on a timescale considered in Fig. 1M: *D*_*x*_ = *D*_*x*_ = 0.8 and *D*_*θ*_ = 0.4. We refer to this model as Model 0.

Equations (1a,b) were solved by using the Euler-Maruyama method with timestep of 0.001. The typical dynamics of the particle for *f* = 0 and *f* = 1 are illustrated in Fig. S2B and S2C, respectively. When *f* = 0, the angle *θ* increases constantly (Fig. S2C, the top panel). As a result, the length of the time during which the particle moves upward is almost equal to the downward counterpart. In contrast, when *f* = 1, the particle tends to spend a longer time around *θ* = *π*/2 (Fig. S2C, the top panel), resulting in larger *y* values than those for *f* = 0 (Fig. S2B and S2C, the bottom panels). Replacing *v*(*θ*; 0) and *ω*(*θ*; 0), which are both constant in Model 0, with experimental data under the no-flow condition does not change the result qualitatively. We conducted simulation of a model where *v*(*θ*; 0) and *ω*(*θ*; 0) are set in the same way as *v*(*θ*; 1) and *ω*(*θ*; 1) except that we use the experimental data from 0 to 1 min, during which flow stimulation is not applied. A representative trajectory, which is illustrated in Fig. S2D, wanders around the initial position drawing helical shape. Furthermore, like the case of Model 0 with *f* = 0, the slope of the MSD is close to 1/2, indicating diffusive dynamics (Fig. S2E).

To break down the contributions to the rheotaxis from the modulation of the speed and angular velocity, we have conducted “one-sided” simulations as below. First, we imposed the heading-speed-only condition, where we replaced *ω*(*θ*; 1) with *ω*_0_ in Model 0 and set *f* = 1. The MSD under this condition, plotted with the solid green curve in Fig. S2E, indicates almost diffusive dynamics in sufficiently large timescales. Next, we performed simulation under the angular-velocity-only condition, where we replaced *v*(*θ*; 1) with *v*_0_ in Model 0 and set *f* = 1. In contrast to the heading-speed-only condition, MSD under the angular-velocity-only condition has a larger slope than that of Model 0 (Orenge solid curve in Fig. S2E). This implies that the modulation of the angular velocity according to the body orientation angle is the primary dynamical mechanism of rheotaxis.

Is the modulation of the heading speed irrelevant to rheotaxis? The following observation implies that the answer is no. Of the two troughs in the plot of heading speed under the flow condition (Fig. S4H), the one around *θ* = 90° can be, at least to some extent, explained as a passive response to the flow because the flow can cancel out the force that *Ciona* generates to swim forward. In contrast, the trough around *θ* = 270° can not be explained by such compensation. To investigate how the latter trough affects rheotaxis, we conducted numerical simulation under the condition where *f* = 1 and the trough of *v*(*θ*; 1) around *θ* = 270° is removed. More specifically, for *π* ≤ *θ* < 2 *π*, we replace *v*(*θ*; 1) in Model 0 with a constant *v*_1_, where *v*_1_ is the mean of the heading speed over the body orientation angle from 180° to 360° in the experiments under the flow condition. Then, the simulation of this model for 1000 [unit time] was performed with 100 different noise realizations. The mean displacement in the y direction from the initial position averaged over 100 realizations was -8.9 [unit length], while that of Model 0 was 18.5. Therefore, the trough around *θ* = 270° is considered to enhance rheotaxis.

## REFERENCES

1. Alerstam, T., Hedenström, A., and Åkesson, S. (2003). Long-distance migration: evolution and determinants. Oikos 103, 247–260. 10.1034/j.1600-0706.2003.12559.x.

2. Dingle, H., and Drake, V.A. (2007). What is migration? Bioscience 57, 113–121.

3. Gibson, B.M., Furbish, D.J., Rahman, I.A., Schmeeckle, M.W., Laflamme, M., and Darroch, S.A.F. (2021). Ancient life and moving fluids. Biol Rev Camb Philos Soc 96, 129–152. 10.1111/brv.12649.

4. Gazzola, M., Argentina, M., and Mahadevan, L. (2014). Scaling macroscopic aquatic locomotion. Nature Physics 10, 758–761. 10.1038/nphys3078.

5. Dudley, R. (2000). The Biomechanics of Insect Flight Form, Function, Evolution (Princeton University Press).

6. Stupski, S.D., and van Breugel, F. (2024). Wind gates olfaction-driven search states in free flight. Curr Biol. 10.1016/j.cub.2024.07.009.

7. Chapman, J.W., Klaassen, R.H., Drake, V.A., Fossette, S., Hays, G.C., Metcalfe, J.D., Reynolds, A.M., Reynolds, D.R., and Alerstam, T. (2011). Animal orientation strategies for movement in flows. Curr Biol 21, R861–870. 10.1016/j.cub.2011.08.014.

8. Paterson, J.R., Gehling, J.G., Droser, M.L., and Bicknell, R.D. (2017). Rheotaxis in the Ediacaran epibenthic organism Parvancorina from South Australia. Sci Rep 7, 45539. 10.1038/srep45539.

9. Darroch, S.A.F., Rahman, I.A., Gibson, B., Racicot, R.A., and Laflamme, M. (2017). Inference of facultative mobility in the enigmatic Ediacaran organism Parvancorina. Biol Lett 13. 10.1098/rsbl.2017.0033.

10. Coutts, F.J., Bradshaw, C.J.A., García-Bellido, D.C., and Gehling, J.G. (2018). Evidence of sensory-driven behavior in the Ediacaran organism Parvancorina: Implications and autecological interpretations. Gondwana Research 55, 21–29. 10.1016/j.gr.2017.10.009.

11. Lyon, E.P. (1904). ON RHEOTROPISM. I. — RHEOTROPISM IN FISHES. American Journal of Physiology-Legacy Content 12, 149–161. 10.1152/ajplegacy.1904.12.2.149.

12. Mathewson, R.F., and Hodgson, E.S. (1972). Klinotaxis and rheotaxis in orientation of sharks toward chemical stimuli. Comparative Biochemistry and Physiology Part A: Physiology 42, 79–84. 10.1016/0300-9629(72)90369-6.

13. Montgomery, J.C., Baker, C.F., and Carton, A.G. (1997). The lateral line can mediate rheotaxis in fish. Nature 389, 960–963. 10.1038/40135.

14. Baker, C.F., and Montgomery, J.C. (1999). The sensory basis of rheotaxis in the blind Mexican cave fish, Astyanax fasciatus. Journal of Comparative Physiology A 184, 519–527. 10.1007/s003590050351.

15. Gardiner, J.M., and Atema, J. (2007). Sharks need the lateral line to locate odor sources: rheotaxis and eddy chemotaxis. Journal of Experimental Biology 210, 1925–1934. 10.1242/jeb.000075.

16. Suli, A., Watson, G.M., Rubel, E.W., and Raible, D.W. (2012). Rheotaxis in larval zebrafish is mediated by lateral line mechanosensory hair cells. PLoS One 7, e29727. 10.1371/journal.pone.0029727.

17. Olive, R., Wolf, S., Dubreuil, A., Bormuth, V., Debregeas, G., and Candelier, R. (2016). Rheotaxis of Larval Zebrafish: Behavioral Study of a Multi-Sensory Process. Front Syst Neurosci 10, 14. 10.3389/fnsys.2016.00014.

18. Oteiza, P., Odstrcil, I., Lauder, G., Portugues, R., and Engert, F. (2017). A novel mechanism for mechanosensory-based rheotaxis in larval zebrafish. Nature 547, 445-448 10.1038/nature23014.

19. Durand, J.P., and Parzefall, J. (1987). Comparative study of the rheotaxis in the cave salamander Proteus anguinus and his epigean relative Necturus maculosus (Proteidae, Urodela). Behavioural Processes 15, 285–291. 10.1016/0376-6357(87)90013-1.

20. Kobayashi, D.R., Farman, R., Polovina, J.J., Parker, D.M., Rice, M., and Balazs, G.H. (2014). “Going with the flow” or not: evidence of positive rheotaxis in oceanic juvenile loggerhead turtles (Caretta caretta) in the South Pacific Ocean Using Satellite Tags and Ocean Circulation Data. PLoS One 9, e103701. 10.1371/journal.pone.0103701.

21. Rowat, D., and Gore, M. (2007). Regional scale horizontal and local scale vertical movements of whale sharks in the Indian Ocean off Seychelles. Fisheries Research 84, 32–40. 10.1016/j.fishres.2006.11.009.

22. Orger, M.B., and Baier, H. (2005). Channeling of red and green cone inputs to the zebrafish optomotor response. Vis Neurosci 22, 275–281. 10.1017/S0952523805223039.

23. Orger, M.B., Smear, M.C., Anstis, S.M., and Baier, H. (2000). Perception of Fourier and non-Fourier motion by larval zebrafish. Nature Neuroscience 3, 1128-1133.10.1038/80649.

24. Orger, M.B., Kampff, A.R., Severi, K.E., Bollmann, J.H., and Engert, F. (2008). Control of visually guided behavior by distinct populations of spinal projection neurons. Nature Neuroscience 11, 327–333. 10.1038/nn2048.

25. Arnold, G.P. (1974). RHEOTROPISM IN FISHES. Biological Reviews 49, 515–576. 10.1111/j.1469-185X.1974.tb01173.x.

26. Bak-Coleman, J., Smith, D., and Coombs, S. (2015). Going with, then against the flow: evidence against the optomotor hypothesis of fish rheotaxis. Animal Behaviour 107, 7–17. 10.1016/j.anbehav.2015.06.007.

27. Valera, G., Markov, D.A., Bijari, K., Randlett, O., Asgharsharghi, A., Baudoin, J.P., Ascoli, G.A., Portugues, R., and Lopez-Schier, H. (2021). A neuronal blueprint for directional mechanosensation in larval zebrafish. Curr Biol 31, 1463–1475 e1466. 10.1016/j.cub.2021.01.045.

28. Chagnaud, B.P., Brucker, C., Hofmann, M.H., and Bleckmann, H. (2008). Measuring flow velocity and flow direction by spatial and temporal analysis of flow fluctuations. J Neurosci 28, 4479–4487. 10.1523/JNEUROSCI.4959-07.2008.

29. Kulpa, M., Bak-Coleman, J., and Coombs, S. (2015). The lateral line is necessary for blind cavefish rheotaxis in non-uniform flow. J Exp Biol 218, 1603–1612. 10.1242/jeb.119537.

30. Engelmann, J., Hanke, W., Mogdans, J., and Bleckmann, H. (2000). Hydrodynamic stimuli and the fish lateral line. Nature 408, 51–52. 10.1038/35040706.

31. Coombs, S., Bak-Coleman, J., and Montgomery, J. (2020). Rheotaxis revisited: a multi-behavioral and multisensory perspective on how fish orient to flow. J Exp Biol 223. 10.1242/jeb.223008.

32. Vanwalleghem, G., Schuster, K., Taylor, M.A., Favre-Bulle, I.A., and Scott, E.K. (2020). Brain-Wide Mapping of Water Flow Perception in Zebrafish. J Neurosci 40, 4130–4144. 10.1523/JNEUROSCI.0049-20.2020.

33. Visser, A.W. (2001). Hydromechanical signals in the plankton. Marine Ecology Progress Series 222, 1-24. DOI 10.3354/meps222001.

34. Hamner, W.M., Hamner, P.P., Strand, S.W., and Gilmer, R.W. (1983). Behavior of Antarctic Krill, Euphausia superba: Chemoreception, Feeding, Schooling, and Molting. Science 220, 433–435. 10.1126/science.220.4595.433.

35. Fossette, S., Gleiss, A.C., Chalumeau, J., Bastian, T., Armstrong, C.D., Vandenabeele, S., Karpytchev, M., and Hays, G.C. (2015). Current-oriented swimming by jellyfish and its role in bloom maintenance. Curr Biol 25, 342–347. 10.1016/j.cub.2014.11.050.

36. McManus, M.A., and Woodson, C.B. (2012). Plankton distribution and ocean dispersal. J Exp Biol 215, 1008–1016. 10.1242/jeb.059014.

37. Genin, A., Jaffe, J.S., Reef, R., Richter, C., and Franks, P.J.S. (2005). Swimming Against the Flow: A Mechanism of Zooplankton Aggregation. Science 308, 860–862. doi:10.1126/science.1107834.

38. Michalec, F.G., Fouxon, I., Souissi, S., and Holzner, M. (2017). Zooplankton can actively adjust their motility to turbulent flow. Proc Natl Acad Sci U S A 114, E11199–E11207. 10.1073/pnas.1708888114.

39. Fay, R.C., and Johnson, J.V. (1971). Observations on the Distribution and Ecology of the Littoral Ascidians of the Mainland Coast of Southern California. Bulletin, Southern California Academy of Sciences 70, 114–124.

40. Hernández-Zanuy, A., and Carballo, J. (2001). Distribution and abundance of ascidian assemblages in Caribbean reef zones of the Golfo de Batabanó (Cuba). Coral Reefs 20, 159–162. 10.1007/s003380100154.

41. Lambert, C.C., and Lambert, G. (2003). Persistence and differential distribution of nonindigenous ascidians in harbors of the Southern California Bight. Marine Ecology Progress Series 259, 145–161.

42. Ryan, K., and Meinertzhagen, I.A. (2019). Neuronal identity: the neuron types of a simple chordate sibling, the tadpole larva of Ciona intestinalis. Curr Opin Neurobiol 56, 47–60. 10.1016/j.conb.2018.10.015.

43. Nakagawa, M., Miyamoto, T., Ohkuma, M., and Tsuda, M. (1999). Action Spectrum for the Photophobic Response of Ciona intestinalis (Ascidieacea, Urochordata) Larvae Implicates Retinal Protein. Photochemistry and Photobiology 70, 359–362.

44. Kajiwara, S., and Yoshida, M. (1985). Changes in Behavior and Ocellar Structure during the Larval Life of Solitary Ascidians. Biol Bull 169, 565-577. Doi 10.2307/1541299.

45. Zega, G., Thorndyke, M.C., and Brown, E.R. (2006). Development of swimming behaviour in the larva of the ascidian Ciona intestinalis. J Exp Biol 209, 3405–3412. 10.1242/jeb.02421.

46. Young, C.M., and Chia, F.-S. (1985). An experimental test of shadow response function in ascidian tadpoles. Journal of Experimental Marine Biology and Ecology 85, 165–175. 10.1016/0022-0981(85)90141-8.

47. Mathis, A., Mamidanna, P., Cury, K.M., Abe, T., Murthy, V.N., Mathis, M.W., and Bethge, M. (2018). DeepLabCut: markerless pose estimation of user-defined body parts with deep learning. Nat Neurosci 21, 1281–1289. 10.1038/s41593-018-0209-y.

48. Athira, A., Dondorp, D., Rudolf, J., Peytral, O., and Chatzigeorgiou, M. (2022). Comprehensive analysis of locomotion dynamics in the protochordate Ciona intestinalis reveals how neuromodulators flexibly shape its behavioral repertoire. PLoS Biol 20, e3001744. 10.1371/journal.pbio.3001744.

49. Hiraiwa, T. (2019). Two types of exclusion interactions for self-propelled objects and collective motion induced by their combination. Phys Rev E 99, 012614. 10.1103/PhysRevE.99.012614.

50. Ryan, K., Lu, Z., and Meinertzhagen, I.A. (2017). The peripheral nervous system of the ascidian tadpole larva: Types of neurons and their synaptic networks. J Comp Neurol. 10.1002/cne.24353.

51. Konno, A., Kaizu, M., Hotta, K., Horie, T., Sasakura, Y., Ikeo, K., and Inaba, K. (2010). Distribution and structural diversity of cilia in tadpole larvae of the ascidian Ciona intestinalis. Dev Biol 337, 42–62. 10.1016/j.ydbio.2009.10.012.

52. Terakubo, H.Q., Nakajima, Y., Sasakura, Y., Horie, T., Konno, A., Takahashi, H., Inaba, K., Hotta, K., and Oka, K. (2010). Network structure of projections extending from peripheral neurons in the tunic of ascidian larva. Dev Dyn 239, 2278–2287. 10.1002/dvdy.22361.

53. Anselmi, C., Fuller, G.K., Stolfi, A., Groves, A.K., and Manni, L. (2024). Sensory cells in tunicates: insights into mechanoreceptor evolution. Front Cell Dev Biol 12, 1359207. 10.3389/fcell.2024.1359207.

54. Hoyer, J., Kolar, K., Athira, A., van den Burgh, M., Dondorp, D., Liang, Z., and Chatzigeorgiou, M. (2024). Polymodal sensory perception drives settlement and metamorphosis of Ciona larvae. Curr Biol 34, 1168–1182 e1167. 10.1016/j.cub.2024.01.041.

55. Poncelet, G., Parolini, L., and Shimeld, S.M. (2024). A microfluidic chip for immobilization and imaging of Ciona intestinalis larvae. J Exp Zool B Mol Dev Evol. 10.1002/jez.b.23267.

56. Wakai, M.K., Nakamura, M.J., Sawai, S., Hotta, K., and Oka, K. (2021). Two-Round Ca(2+) transient in papillae by mechanical stimulation induces metamorphosis in the ascidian Ciona intestinalis type A. Proc Biol Sci 288, 20203207. 10.1098/rspb.2020.3207.

57. Sakamoto, A., Hozumi, A., Shiraishi, A., Satake, H., Horie, T., and Sasakura, Y. (2022). The TRP channel PKD2 is involved in sensing the mechanical stimulus of adhesion for initiating metamorphosis in the chordate Ciona. Dev Growth Differ 64, 395–408. 10.1111/dgd.12801.

58. Chen, T.W., Wardill, T.J., Sun, Y., Pulver, S.R., Renninger, S.L., Baohan, A., Schreiter, E.R., Kerr, R.A., Orger, M.B., Jayaraman, V., et al. (2013). Ultrasensitive fluorescent proteins for imaging neuronal activity. Nature 499, 295–300. 10.1038/nature12354.

59. Kolar, K., Dondorp, D., Zwiggelaar, J.C., Høyer, J., and Chatzigeorgiou, M. (2021). Mesmerize is a dynamically adaptable user-friendly analysis platform for 2D and 3D calcium imaging data. Nature Communications 12. 10.1038/s41467-021-26550-y.

60. Ryan, K., Lu, Z., and Meinertzhagen, I.A. (2016). The CNS connectome of a tadpole larva of Ciona intestinalis (L.) highlights sidedness in the brain of a chordate sibling. Elife 5. 10.7554/eLife.16962.

61. Khan, N.A., Willemarck, N., Talebi, A., Marchand, A., Binda, M.M., Dehairs, J., Rueda-Rincon, N., Daniels, V.W., Bagadi, M., Thimiri Govinda Raj, D.B., et al. (2016). Identification of drugs that restore primary cilium expression in cancer cells. Oncotarget 7, 9975–9992. 10.18632/oncotarget.7198.

62. Buskey, E.J., Lenz, P.H., and Hartline, D.K. (2011). Sensory perception, neurobiology, and behavioral adaptations for predator avoidance in planktonic copepods. Adaptive Behavior 20, 57–66. 10.1177/1059712311426801.

63. Pigolotti, S., and Benzi, R. (2014). Selective advantage of diffusing faster. Phys Rev Lett 112, 188102. 10.1103/PhysRevLett.112.188102.

64. Zhao, D., Chen, S., Horie, T., Gao, Y., Bao, H., and Liu, X. (2020). Comparison of differentiation gene batteries for migratory mechanosensory neurons across bilaterians. Evol Dev 22, 438–450. 10.1111/ede.12331.

65. Pasini, A., Amiel, A., Rothbacher, U., Roure, A., Lemaire, P., and Darras, S. (2006). Formation of the ascidian epidermal sensory neurons: insights into the origin of the chordate peripheral nervous system. PLoS Biol 4, e225. 10.1371/journal.pbio.0040225.

66. Desban, L., Roussel, J., Mirat, O., Lejeune, F.-X., Keiser, L., Michalski, N., and Wyart, C. (2022). Lateral line hair cells integrate mechanical and chemical cues to orient navigation. bioRxiv, 2022.2008.2031.505989. 10.1101/2022.08.31.505989.

67. Mogdans, J., Bleckmann, H., and Menger, N. (2008). Sensitivity of Central Units in the Goldfish, Carassius auratus, to Transient Hydrodynamic Stimuli (Part 1 of 2). Brain Behav Evolut 50, 261–272. 10.1159/000113341.

68. Künzel, S., Bleckmann, H., and Mogdans, J. (2011). Responses of brainstem lateral line units to different stimulus source locations and vibration directions. Journal of Comparative Physiology A 197, 773–787. 10.1007/s00359-011-0642-9.

69. Torrence, S.A., and Cloney, R.A. (1982). Nervous-System of Ascidian Larvae - Caudal Primary Sensory Neurons. Zoomorphology 99, 103-115. Doi 10.1007/Bf00310303.

70. Marques, J.C., Li, M., Schaak, D., Robson, D.N., and Li, J.M. (2020). Internal state dynamics shape brainwide activity and foraging behaviour. Nature 577, 239–243. 10.1038/s41586-019-1858-z.

71. Gordus, A., Pokala, N., Levy, S., Flavell, S.W., and Bargmann, C.I. (2015). Feedback from network states generates variability in a probabilistic olfactory circuit. Cell 161, 215–227. 10.1016/j.cell.2015.02.018.

72. Flavell, S.W., Pokala, N., Macosko, E.Z., Albrecht, D.R., Larsch, J., and Bargmann, C.I. (2013). Serotonin and the neuropeptide PDF initiate and extend opposing behavioral states in C. elegans. Cell 154, 1023–1035. 10.1016/j.cell.2013.08.001.

73. Lottem, E., Banerjee, D., Vertechi, P., Sarra, D., Lohuis, M.O., and Mainen, Z.F. (2018). Activation of serotonin neurons promotes active persistence in a probabilistic foraging task. Nat Commun 9, 1000. 10.1038/s41467-018-03438-y.

74. Kamiya, C., Ohta, N., Ogura, Y., Yoshida, K., Horie, T., Kusakabe, T.G., Satake, H., and Sasakura, Y. (2014). Nonreproductive role of gonadotropin-releasing hormone in the control of ascidian metamorphosis. Dev Dyn 243, 1524–1535. 10.1002/dvdy.24176.

75. Okawa, N., Shimai, K., Ohnishi, K., Ohkura, M., Nakai, J., Horie, T., Kuhara, A., and Kusakabe, T.G. (2020). Cellular identity and Ca(2+) signaling activity of the non-reproductive GnRH system in the Ciona intestinalis type A (Ciona robusta) larva. Sci Rep 10, 18590. 10.1038/s41598-020-75344-7.

76. Horie, T., Nakagawa, M., Sasakura, Y., Kusakabe, T.G., and Tsuda, M. (2010). Simple motor system of the ascidian larva: neuronal complex comprising putative cholinergic and GABAergic/glycinergic neurons. Zoolog Sci 27, 181–190. 10.2108/zsj.27.181.

77. Horie, T., Kusakabe, T., and Tsuda, M. (2008). Glutamatergic networks in the Ciona intestinalis larva. J Comp Neurol 508, 249–263. 10.1002/cne.21678.

78. Takamura, K., Minamida, N., and Okabe, S. (2010). Neural map of the larval central nervous system in the ascidian Ciona intestinalis. Zoolog Sci 27, 191–203. 10.2108/zsj.27.191.

79. Brown, E.R., Nishino, A., Bone, Q., Meinertzhagen, I.A., and Okamura, Y. (2005). GABAergic synaptic transmission modulates swimming in the ascidian larva. Eur J Neurosci 22, 2541–2548. 10.1111/j.1460-9568.2005.04420.x.

80. Zega, G., Biggiogero, M., Groppelli, S., Candiani, S., Oliveri, D., Parodi, M., Pestarino, M., De Bernardi, F., and Pennati, R. (2008). Developmental expression of glutamic acid decarboxylase and of gamma-aminobutyric acid type B receptors in the ascidian Ciona intestinalis. J Comp Neurol 506, 489–505. 10.1002/cne.21565.

81. Schindelin, J., Arganda-Carreras, I., Frise, E., Kaynig, V., Longair, M., Pietzsch, T., Preibisch, S., Rueden, C., Saalfeld, S., Schmid, B., et al. (2012). Fiji: an open-source platform for biological-image analysis. Nat Methods 9, 676–682. 10.1038/nmeth.2019.

82. Rudolf, J., Dondorp, D., Canon, L., Tieo, S., and Chatzigeorgiou, M. (2019). Automated behavioural analysis reveals the basic behavioural repertoire of the urochordate Ciona intestinalis. Sci Rep 9, 2416. 10.1038/s41598-019-38791-5.

83. Liang, Z., Dondorp, D.C., and Chatzigeorgiou, M. (2024). The ion channel Anoctamin 10/TMEM16K coordinates organ morphogenesis across scales in the urochordate notochord. PLoS Biol 22, e3002762. 10.1371/journal.pbio.3002762.

84. Pnevmatikakis, E.A., and Giovannucci, A. (2017). NoRMCorre: An online algorithm for piecewise rigid motion correction of calcium imaging data. J Neurosci Methods 291, 83–94. 10.1016/j.jneumeth.2017.07.031.

85. Pnevmatikakis, E.A., Soudry, D., Gao, Y., Machado, T.A., Merel, J., Pfau, D., Reardon, T., Mu, Y., Lacefield, C., Yang, W., et al. (2016). Simultaneous Denoising, Deconvolution, and Demixing of Calcium Imaging Data. Neuron 89, 285–299. 10.1016/j.neuron.2015.11.037.

86. Giovannucci, A., Friedrich, J., Gunn, P., Kalfon, J., Brown, B.L., Koay, S.A., Taxidis, J., Najafi, F., Gauthier, J.L., Zhou, P., et al. (2019). CaImAn an open source tool for scalable calcium imaging data analysis. Elife 8. 10.7554/eLife.38173.

87. Pedregosa, F., Varoquaux, G., Gramfort, A., Michel, V., Thirion, B., Grisel, O., Blondel, M., Prettenhofer, P., Weiss, R., Dubourg, V., et al. (2011). Scikit-learn: Machine Learning in Python. J Mach Learn Res 12, 2825–2830.

88. Leggio, B., Laussu, J., Carlier, A., Godin, C., Lemaire, P., and Faure, E. (2019). MorphoNet: an interactive online morphological browser to explore complex multi-scale data. Nat Commun 10, 2812. 10.1038/s41467-019-10668-1.

89. Toschi, F., and Sega, M. (2019). Flowing Matter 10.1007/978-3-030-23370-9.

90. van Teeffelen, S., and Löwen, H. (2008). Dynamics of a Brownian circle swimmer. Physical Review E 78, 020101. 10.1103/PhysRevE.78.020101.

91. Nevel’son, M.B. (1964). On the Behavior of the Invariant Measure of a Diffusion Process with Small Diffusion on a Circle. Theory of Probability & Its Applications 9, 125–131. 10.1137/1109016.90.

