## Supplementary material for "The cellular and behavioral blueprints of chordate rheotaxis": Document S1

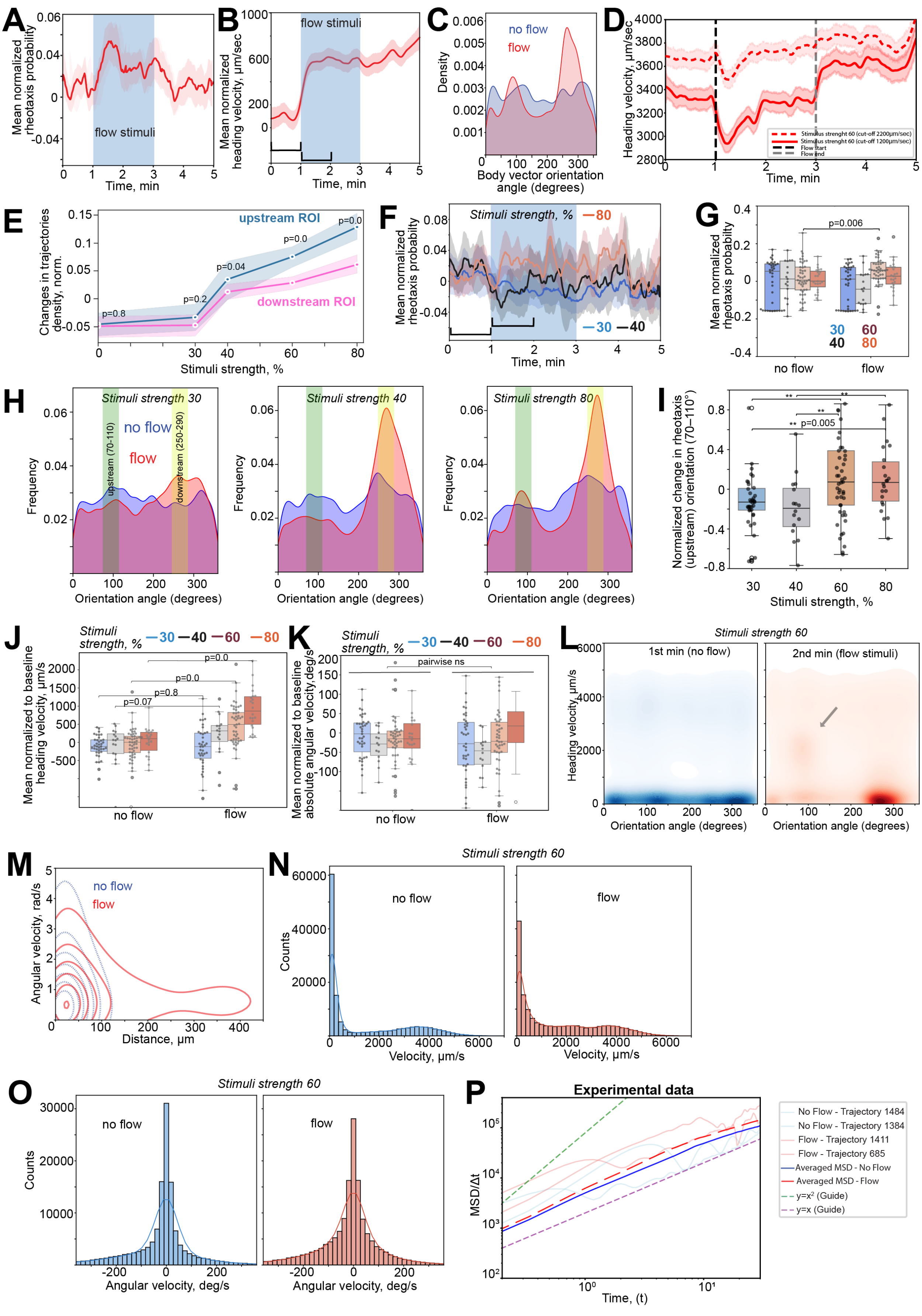

A
Mean normalized rheotaxis probability
flow stimuli
Time, min
B
Mean normalized heading velocity, µm/sec
flow stimuli
Time, min
C
Density
no flow
flow
Body vector orientation angle (degrees)
D
Heading velocity, µm/sec
Time, min
Stimulus strenght 60 (cut-off 2200µm/sec)
Stimulus strenght 60 (cut-off 1200µm/sec)
Flow start
Flow end
E
Changes in trajectories density, norm.
upstream ROI
downstream ROI
p=0.8
p=0.2
p=0.04
p=0.0
p=0.0
Stimuli strength, %
F
Mean normalized rheotaxis probability
Stimuli strength, %
—80
—30 —40
Time, min
G
Mean normalized rheotaxis probability
p=0.006
30 60
40 80
no flow
flow
H
Stimuli strength 30
Stimuli strength 40
Stimuli strength 80
no flow
flow
upstream (70-110)
downstream (250-290)
Frequency
Orientation angle (degrees)
I
Normalized change in rheotaxis (upstream) orientation (70–110°)
p=0.005
Stimuli strength, %
J
Stimuli strength, %
—30 —40 —60 —80
Mean normalized to baseline heading velocity, µm/s
p=0.0
p=0.0
p=0.8
p=0.07
no flow
flow
K
Stimuli strength, %
—30 —40 —60 —80
Mean normalized to baseline absolute angular velocity, deg/s
pairwise ns
no flow
flow
L
Stimuli strength 60
1st min (no flow)
2nd min (flow stimuli)
Heading velocity, µm/s
Orientation angle (degrees)
M
Angular velocity, rad/s
no flow
flow
Distance, µm
N
Stimuli strength 60
no flow
flow
Counts
Velocity, µm/s
O
Stimuli strength 60
no flow
flow
Counts
Angular velocity, deg/s
P
Experimental data
MSD/Δt
Time, (t)
No Flow - Trajectory 1484
No Flow - Trajectory 1384
Flow - Trajectory 1411
Flow - Trajectory 685
Averaged MSD - No Flow
Averaged MSD - Flow
y=x² (Guide)
y=x (Guide)

**Figure S1. Dose-dependent or threshold-like response of rheotaxis to increasing flow strength. Related to Figure 1.**

**(A)** Mean rheotaxis probability under no flow conditions and 60% strength, i.e. 2 mm/s. Solid line shows the mean probability value, shade indicates SEM. Blue vertical bar defines the flow stimulus presentation for the flow experiments (same for panel B). **(B)** Mean normalized heading speed for 2mm/s flow. **(C)** Body vector orientation distributions under no flow (blue) and 60% flow (red) conditions for wild-type larvae. **(D)** Heading speed for 2 mm/s flow with data filtered to include animal speed fragments > 2.2 mm/s (dashed line), >1.2 mm/s (solid line) to minimize impact of animals potentially dragged downstream by the flow. **(E)** Quantification of trajectory density reveals a dose-dependent increase in upstream densities in response to different flow strengths compared to the control condition (no stimulus). Stimulus strengths are defined as follows: 30% = 0.5 mm/s (0.5 body length/s), 40% = 1 mm/sec (1 body length/s), 60% = 2 mm/s (2 body lengths/s), and 80% = 3 mm/s (3 body lengths/s). **(F)** Rheotaxis probability at different stimulus strengths, quantified using a pose estimation approach. Note that 60% stimuli mean rheotaxis probability is present in Fig 1H. Mean rheotaxis probabilities were normalized to baseline (the median of the no-flow condition) for each experiment. Bars below indicate the time intervals used for quantification. **(G)** Quantification of mean rheotaxis probability across different stimulus strengths. **(H)** Orientation angles distribution for different stimuli strengths with highlighted bins of up and downstream angles. Note, stimuli 60 is presented in the Figure 1I. **(I)** Quantification of the change in rheotaxis (upstream) orientation angles distribution normalized to the distribution in no flow condition for each experiment. **(J)** Mean heading speed quantification shows stimulus-dependent changes, indicating a dose-dependent response to counteracting flow. Heading speeds were normalized to baseline (the first second of the no-flow condition) for each experiment. **(K)** Mean absolute angular velocity quantification shows no significant change across different stimulus strengths. Data were normalized to baseline (the first second of the no-flow condition) for each experiment. **(L)** Kernel density estimate (KDE) plots for heading speed and orientation angles for 60 % stimuli. Arrow = cluster of high-speed upstream swimmers, **(M)** Contour plots showing the quantification of angular velocity as a function of swimming distance, contours represent the KDE density clusters with different distribution between no flow and flow conditions. **(N)** Distribution histograms of heading speed for 60% stimulus strength. **(O)** Distribution histograms of angular velocity for 60% stimulus strength. Data points in G, I, J, K represent individual experiments. Statistical analysis in (H and I) was performed using the Mann-Whitney U test to compare no-flow and flow conditions for each stimulus strength. **(P)** Mean square displacement (MSD) based on experimental data without filtering under no flow (Blue solid curve for averaged MSD and light blue for representative single animals MSDs) or flow stimulus (Red dotted curve for averaged MSD and light solid curves for representative single animals MSDs) conditions showing a shift (transition) of the movement pattern from a more diffusive regime (purple guideline) characterized by frequent reorientations to a more ballistic regime (green guideline) with more directed trajectories in the flow condition.

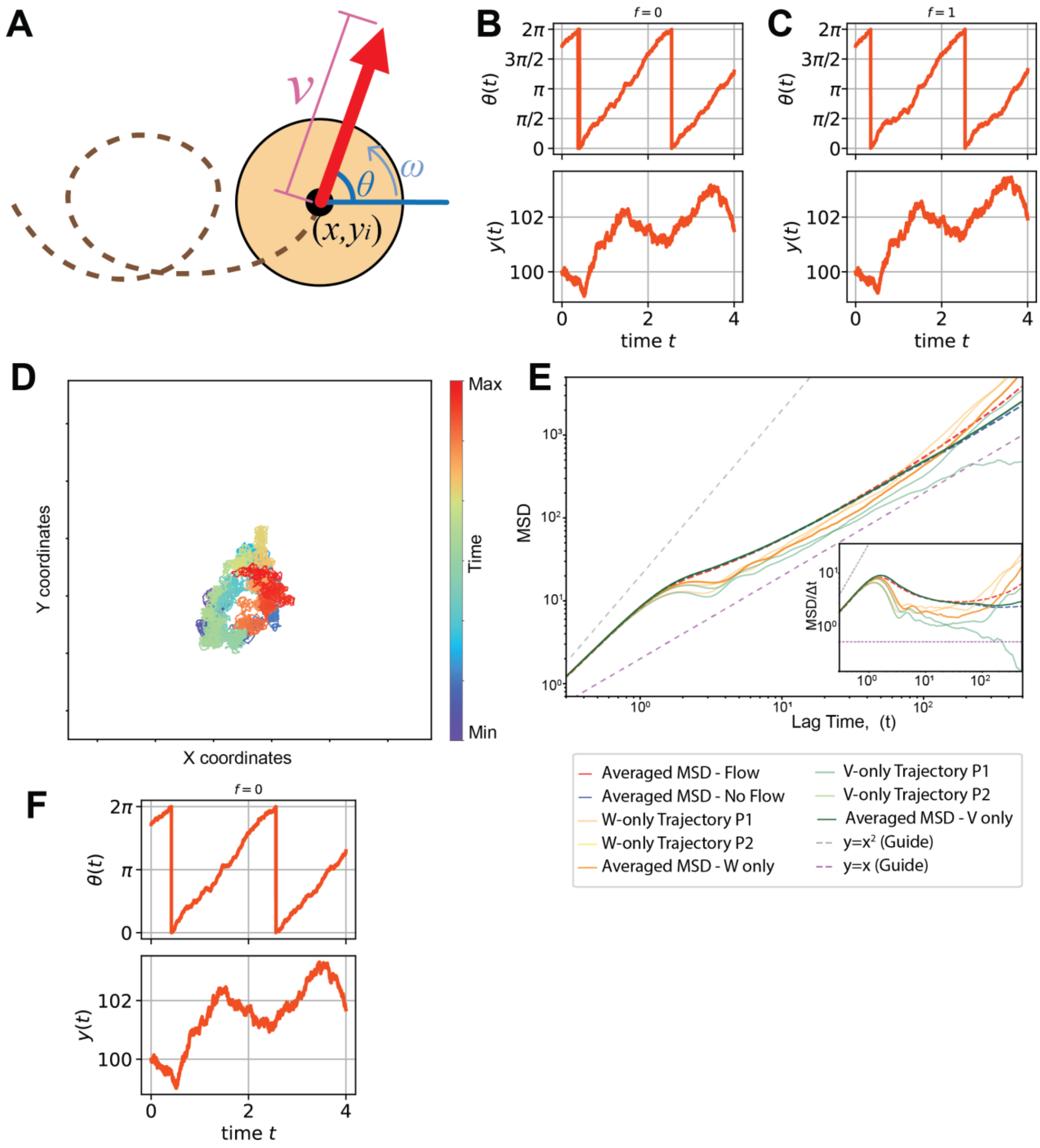


**Figure S2. Mathematical models of the swimming trajectory of *Ciona* with and without flow. Related to Figure 1.**

**(A)** Schematic illustrating the heading speed and angular velocity vectors of swimming *Ciona* larvae used in experimental analysis and mathematical modelling. **(B, C)** Typical time series of the angle $\theta$ and $y$ coordinate of the model defined by Eq. (1a) and (1b). In the absence of flow ($f = 0$), the angle increases almost constantly apart from fluctuations due to noise (the upper panel of (**A**)). In contrast, in the presence of the flow ($f = 1$), the angle $\theta$ increases slowly at around $\pi/2$ (the upper panel of (**C**)), which causes the particle to spend more time moving upward and hence increased $y$ values (the lower panel of (**C**)) compared with the case of $f = 0$ (the lower panel of (**B**)). The same seed of the random number generator is used for both **(B)** and (**C**). (**D, E, F**) The dynamics of the mathematical model based on the data collected in the one-minute period preceding the onset of the flow stimulation. Like the case of Fig. 1M (left panel), the trajectory (**D**) does not show any tendency to move in a particular direction. Consistently, the MSD (**E**) indicates diffusive dynamics in a sufficiently long timescale. The similarity to the model with constant $v$ and $\omega$ is also observed in the time series of angle $\theta$ and $y$ coordinate (**F**). Note that the angle $\theta$ increases almost constantly as in (**B**). The seed of the random number generator is the same as (**B**) and (**C**).

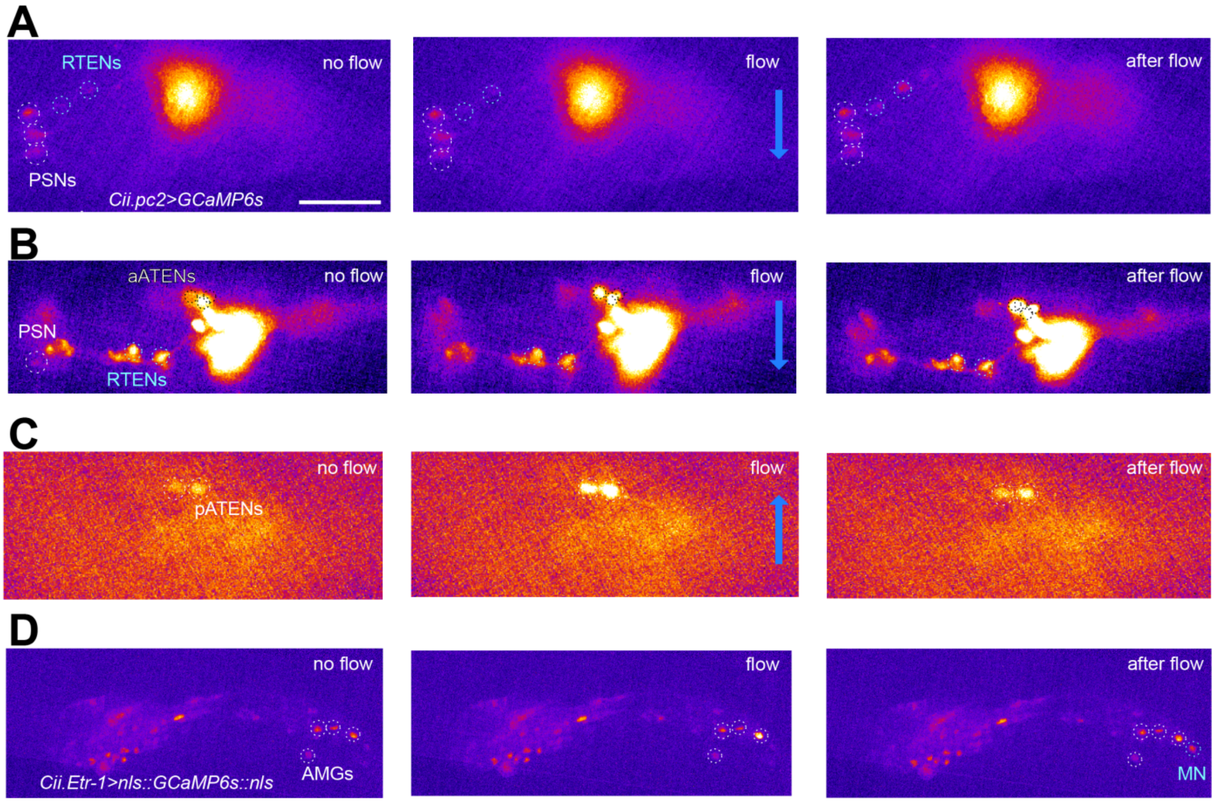


**Figure S3. Directional flow of sea water elicits calcium responses in the larval brain. Related to Figure 2.**

**(A-C)** Snapshots of the trunk region of the representative transgenic *Ciona* larva expressing *Cii.pc2>GCaMP6s* in peptidergic neurons of the CNS and Peripheral Nervous System (PNS) under no-flow and flow conditions. Note the polymodal papillae sensory neurons (PSNs), the rostral trunk epidermal neurons (RTENs), the anterior/ posterior apical trunk epidermal neurons (aATENs & pATENs). Sensory neurons of the same type response differently depending of flow orientation. **(D)** Snapshots of the trunk region of a representative transgenic *Ciona* larva which was used for whole-brain imaging expressing *Cii.Etr-1>nls::GCaMP6s::nls* (green) and *Cii.Etr-1>Lck-mScarlet* (magenda). Note peripheral motor neurons (AMGs) and Motor Neuron (MN) activated by flow stimuli. Scalebar = 50µm.

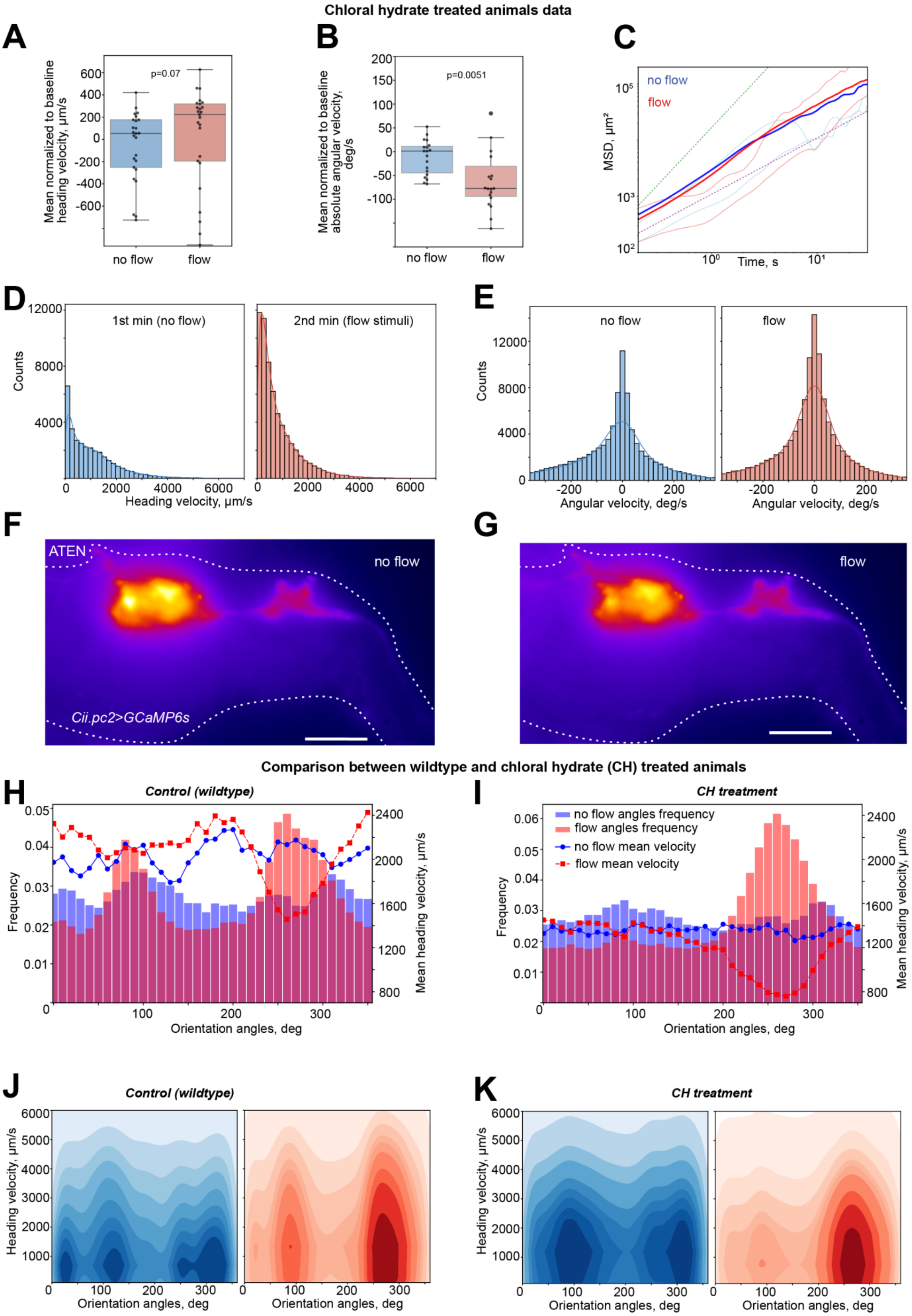
Chloral hydrate treated animals data
A
Mean normalized to baseline heading velocity, µm/s
p=0.07
no flow
flow
B
Mean normalized to baseline absolute angular velocity, deg/s
p=0.0051
no flow
flow
C
MSD, µm²
no flow
flow
Time, s
D
Counts
1st min (no flow)
2nd min (flow stimuli)
Heading velocity, µm/s
E
Counts
no flow
flow
Angular velocity, deg/s
F
ATEN
no flow
Cii.pc2>GCaMP6s
G
flow
Comparison between wildtype and chloral hydrate (CH) treated animals
H
Control (wildtype)
Frequency
Mean heading velocity, µm/s
Orientation angles, deg
I
CH treatment
no flow angles frequency
flow angles frequency
no flow mean velocity
flow mean velocity
Frequency
Mean heading velocity, µm/s
Orientation angles, deg
J
Control (wildtype)
Heading velocity, µm/s
Orientation angles, deg
K
CH treatment
Heading velocity, µm/s
Orientation angles, deg

**Figure S4. Impaired rheotaxis in sensory cilia depleted *Ciona* larvae. Related to Figure 3.**

**(A)** Mean heading speed quantification shows no significant change to the 2 mm/sec flow stimulus. Heading speeds were normalized to baseline (the first second of the no-flow condition) for each experiment. **(B)** Mean absolute angular velocity quantification showed its reduction in response to the flow. Data were normalized to baseline (the first second of the no-flow condition) for each experiment. Data points in a and b represent experiments. Statistical analysis in (A and B) was performed using the Mann-Whitney U test to compare no-flow and flow conditions. **(C)** Mean square displacement (MSD) of CH treated animals under no flow or flow stimulus conditions does not show a prominent shift of the movement pattern from a more diffusive regime (purple guideline) characterized by frequent reorientations to a more ballistic regime (green guideline) with more directed trajectories in the flow condition. **(D)** Distribution histograms of heading speed for chloral hydrate treated animals. **(E)** Distribution histograms of angular velocity for chloral hydrate treated animals. **(G and F)** Snapshots of the trunk region of the representative transgenic *Ciona* larva expressing *Cii.pc2>GCaMP6s* in peptidergic neurons of the CNS and Peripheral Nervous System (PNS) under no-flow and flow conditions. Note the anterior apical trunk epidermal neuron (ATEN) showing no change in calcium activity before and after stimuli application. Scalebar =50 µm. **(H and I)** Relation of orientation angles and the mean heading speed for control and CH treated animals. **(J and K)** Kernel density estimate (KDE) plots of heading speed and orientation angles for control and CH treated animals. For H, I, J and K data was filtered to eliminate lower speed bins < 100 µm per s corresponding to the not moving animals.
